# Cortico-cerebellar beta-band dynamics predict flexible motor timing

**DOI:** 10.64898/2026.09.03.748373

**Authors:** Coen S. Zandvoort, Charlotte C. Wissing, Jonathan Winter, Anna Kowalczyk, Ole Jensen, R. Chris Miall, Devika Narain, Katja Kornysheva

## Abstract

Flexible motor control requires that movements adapt to changing temporal contexts. Here, we test whether flexible timing is driven by context-dependent encoding across cortico-cerebellar circuits and how neural dynamics within these circuits enable accurate performance. To overcome the signal-to-noise limitations of conventional electro- and magnetoencephalography, we recorded whole-head neural dynamics using optically pumped magnetometer arrays (OPM-MEG). Participants learned a context-dependent task, executing manual button presses at time intervals 800 ms (T1) and 1,600 ms (T2), respectively, to avoid a periocular air puff, which was associated with an implicit conditioned eyeblink. We found that the motor cortex contralateral to the hand and bilateral cerebellar lobule VI dynamically encoded these intervals through beta-band (13-30 Hz) event-related desynchronisation (ERD) that precisely scaled with T1 and T2. The cerebellar trial-by-trial latency of the beta-band ERD predicted the timing of explicit manual actions. Finally, partial directed coherence revealed that baseline bidirectional beta-band coupling across the network transiently weakened from the ipsilateral cerebellum to the contralateral motor cortex during finger movement execution. Our findings show that cortico-cerebellar coupling functions as a gating mechanism and suggest that cerebellar circuits modulate cortical motor activity for flexible, accurate motor timing.

**Significance statement:** How human cortico-cerebellar networks flexibly encode context-dependent timing remains poorly understood due to signal-to-noise constraints in non-invasive neural recordings. By combining a task coupling explicit manual and implicit eyeblink responses with wearable optically pumped magnetometers (OPM-MEG), we overcame these depth-sensitivity constraints to resolve human cortico-cerebellar dynamics. Beta-band (13-30 Hz) event-related desynchronisation in the contralateral cortical motor and bilateral cerebellar lobule VI was scaled to the anticipated response timings, with single-trial cerebellar latencies predicting explicit manual response timing. Crucially, directional functional coupling analysis revealed that bidirectional beta-band coupling transiently reduces from cerebellum to motor cortex during movement execution. This approach establishes a blueprint for non-invasively mapping cortico-cerebellar network dynamics, providing a framework to study circuit-level dysfunctions in disorders affecting the cerebellum.

## Introduction

Accurate and flexible movement timing is critical for adaptive behaviour. A long-standing consensus, built upon decades of research, is that temporal control is governed by a widely distributed network of cortical and subcortical structures (Hazeltine et al., 1997; Nobre et al., 2007; Kornysheva, 2016; Remington et al., 2018; Stine and Jazayeri, 2025), yet the underlying distributed neural mechanisms and interaction between these brain regions remain poorly understood (Stine and Jazayeri, 2025). Central to this distributed architecture are cortico-cerebellar loops, comprising closed and reciprocal circuits linking cerebellum and cerebral cortex also encompassing thalamus and brainstem. Recent work in animal models reveals that preparatory cortical motor activity depends fundamentally on cerebellar input to the cortex via the thalamus (Deverett et al., 2018; Gao et al., 2018; Chabrol et al., 2019; Wagner et al., 2019). While non-human primate and rodent models show that flexible motor timing relies on both cerebellar (Koppen et al., 2026) and motor cortical areas (Merchant et al., 2013; Wang et al., 2018), the neural mechanisms coordinating these distributed processes remain unknown.

Recent human behavioural work demonstrates that linking an explicit manual response to an implicit, cerebellar-dependent eyeblink conditioning provides a novel paradigm for probing cortico-cerebellar interactions (Mangili et al., 2025). However, the direct neural mechanisms driving these network interactions have yet to be established. One key impediment to studying the role of the cortico-cerebellar loops in humans has been to obtain cerebellar signals using non-invasive techniques such as electroencephalography (EEG) and magnetoencephalography (MEG) at sufficiently high signal-to-noise ratios (SNR). Whilst cerebellar responses can be captured using non-invasive electrophysiology (Tesche and Karhu, 2000; Kujala et al., 2007), both EEG and MEG suffer from a marked signal-to-noise ratio (SNR) deficit when recording from the cerebellum. Due to the increased distance between cerebellar structures and scalp-surface sensors, cerebellar signal strengths are 30-60% smaller than those recorded from cerebral cortex (Andersen et al., 2020; Samuelsson et al., 2020). However, the factors driving this attenuation differ between modalities: MEG sensitivity is heavily constrained by source orientation and signal cancellation within the convoluted cerebellar folia (Hillebrand and Barnes, 2002; Ahlfors et al., 2010),whereas EEG detects both radial and tangential sources but is primarily limited by volume conduction and signal distortion through the thick suboccipital bone (Nunez and Srinivasan, 2006; Samuelsson et al., 2020). The advent of wearable optically pumped magnetometer (OPM-MEG) arrays enables a closer sensor proximity to and coverage of the cerebellum, and thus improved SNR (Boto et al., 2018; Lin et al., 2019; Lin et al., 2025; West et al., 2025; Roos et al., 2026).

Utilising OPM-MEG we investigated the cortico-cerebellar neural dynamics in humans associated with yoked implicit and explicit motor timing in a task developed by Mangili et al. (2025). Here, participants learnt to produce a flexibly timed manual response based on context signals. Failure to execute a correctly timed manual response resulted in a punitive periocular air puff. Consistent with previous reports (Mangili et al., 2025), in addition to the learning of the context-associated timing of the manual response, a flexibly timed predictive implicit eyeblink response was present on correct trials in which no air puff occurred. Our task design coupled a left-eye air puff with a right-hand manual response to separate the underlying neuroanatomy while examining shared network coordination. While the implicit eyeblink has been linked to ipsilateral left cerebellar lobule VI (Ten Brinke et al., 2015), the right-hand explicit manual response is linked to both left (contralateral) motor cortex and right (ipsilateral) cerebellar lobules IV/V (King et al., 2019). First, we hypothesised that the bilateral cerebellum (Heiney et al., 2014; Ten Brinke et al., 2017; Koppen et al., 2026) and the contralateral motor cortex would encode flexible, context-dependent motor timing (Wang et al., 2018), since the implicit eyeblink and explicit finger press are temporally yoked. Specifically, we predicted that this timing is represented by modulations in the beta band (13-30 Hz) in the motor cortex (Barone and Rossiter, 2021) and the cerebellum (Lin et al., 2025). Second, based on the behavioural evidence suggesting the fast emergence of a context-dependent implicit eyeblink response (Mangili et al., 2025), we hypothesised that implicit-learning-related changes in oscillatory dynamics occur more rapidly in the cerebellum than explicit-learning changes in the cerebral cortex. Third, we hypothesised that cerebellar and cortical motor activity are coupled to enable yoked implicit and explicit learning of the target timing interval.

Our results reveal that both the cerebellum and motor cortex dynamically track context-dependent timing through markers of motor disinhibition – event-related beta-band desynchronisation (beta-band ERD), – which establish more rapidly in cerebellar circuits with training, and correlate with motor timing. Finally, while cerebellar and cortical beta-band ERD covary and exhibit beta-band coupling, single-trial analysis reveals that successful execution is mediated by a well-timed functional weakening of input from the ipsilateral cerebellum to the contralateral motor cortex.

## Materials and methods

### Participants

The study was approved by the University of Birmingham Ethics Committee (under ethics code ERN_18-0226AP34). Fourteen participants were recruited from a student pool which included undergraduate Psychology students as well as volunteering staff from the Centre for Human Brain Health (University of Birmingham). All participants provided written and oral consent throughout the study. Participating Psychology students received credits to compensate for the time spent doing the experiment. Inclusion criteria were (I) being aged between 18 and 35 years of age, (II) having a vision between -1.5 and 1.5 (either uncorrected or corrected using glasses or contact lenses), (III) no formal diagnosis of neurological or neuropsychological disorders (e.g., epilepsy, ADHD), and (IV) having no irremovable parts of metal in their body (e.g., dental work or mechanical implants).

### Experimental design

This study employed the experimental paradigm previously used by Mangili et al. (2025) to study the effects of learning temporal contexts (illustrated in Figure 1). Participants (*n*=14; 8 female and 6 male) were informed about the two different response times depending on the contextual cue that was presented, which aligns to the instructions given to the ‘Strategy’ group (Mangili et al., 2025). These contexts were a dark or light tunnel (Figure 1A), whose image and motion statistics match previous reports (Mangili et al., 2025). We refer to the time interval conditions as T1 and T2, corresponding to dark and light tunnels, respectively. Participants were presented with these stimuli on a screen that was ∼143 cm from their eyes, equalling a visual field span of 19 degrees (ProPixx project, Vpixx Technologies, sampling rate: 120 Hz). Participants completed 450 trials, divided into three blocks of 150 trials. In every trial, the dark or light tunnel (context) trial was decided randomly. After a random interval elapsed after the onset of the context (tunnel), a brief light flash (cue) was presented for 100 ms, after which the tunnel was set in motion at a constant speed while preserving its static visual features that signalled the context. This meant that the end of the tunnel remained far away to minimise the use of depth information during the experiment. After that, the button had to be pressed after 800 ms (omission window: 720-880 ms) in T1 and 1,600 ms (omission window: 1,420-1,760 ms) in T2 for the dark and light tunnel contexts, respectively (Figure 1B). Correctly pressing the button with the right index finger within the designated time interval meant the avoidance of a periocular air puff to the left eye. This punitive, low-latency, gentle, periocular air puff lasted 50 ms and evoked a reflexive eyeblink, akin an unconditioned response. A single air puff was provided on each trial when the button press fell outside the context condition’s time window or after 2,500 ms if no response was made. The context stopped moving after a button press or after 2,500 ms when the participants did not respond. Inter-trial interval was set between 1,000 and 3,000 ms and was randomly sampled from a uniform distribution. The experimental protocol was developed with the Psychophysics Toolbox in MATLAB (version 2019b, MathWorks Inc., Natick, USA) and implemented with VPixx Technologies to create accurate timing.

**Figure 1.**
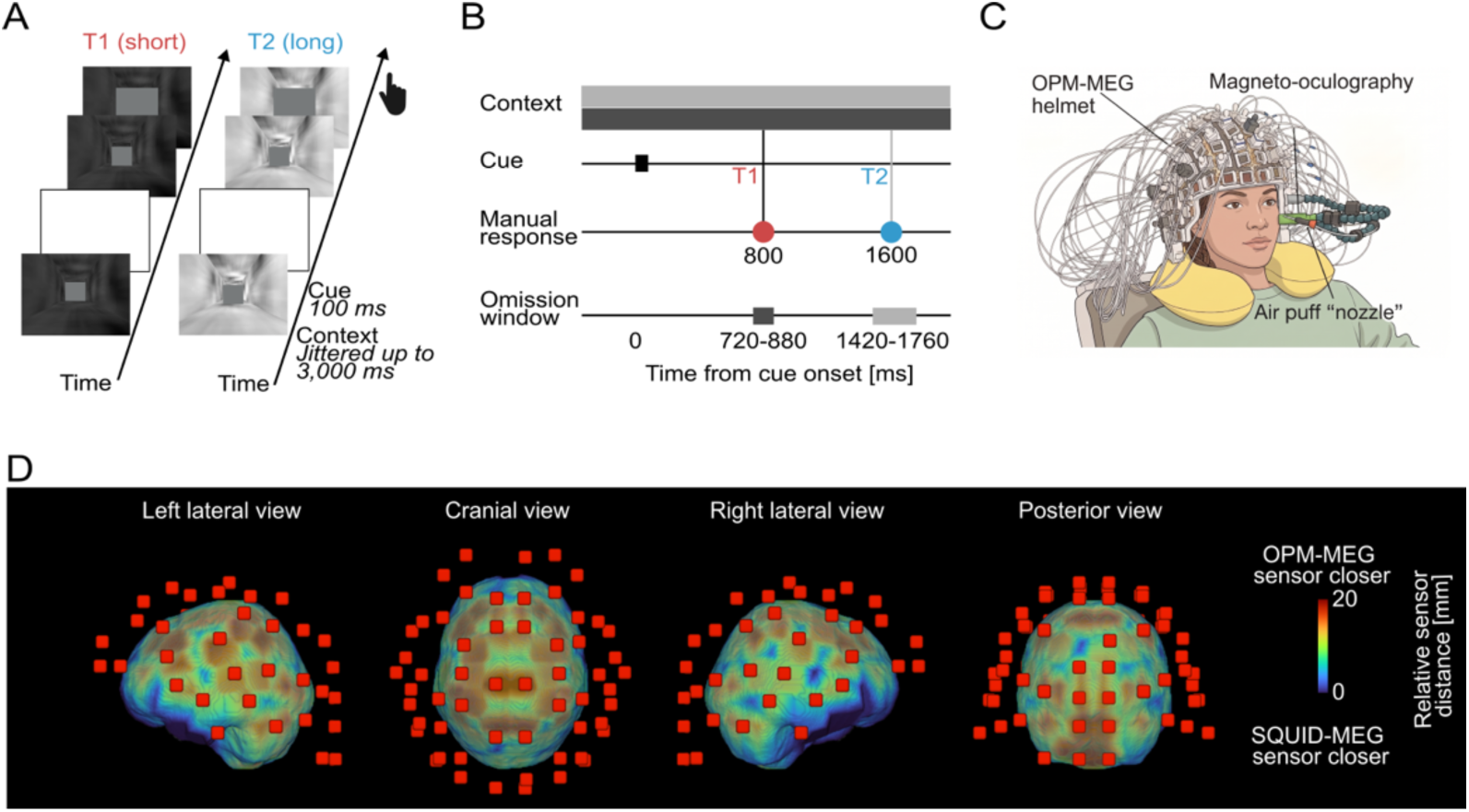
Experimental paradigm and setup. **A)** Trial design with dark and light contexts (referred to as T1 and T2) are presented statically. After a cue (white light flash; 100 ms), the tunnel started moving, after which the participants were meant to correctly press a button with their right index finger after 800 and 1,600 ms. **B)** Participants are presented with two contexts. Participants had to respond after 800 ms (omission: 720-880) and 1,600 (1,420-1,760) for timings T1 and T2, respectively, to avoid an air puff being administered to their left eye. **C)** Participants had an OPM-MEG helmet mounted on their head. Magneto-oculographic (left eye) and magneto-myographic (right trapezius) activity were co-registered. The air-puff nozzle was also integrated in the OPM-MEG helmet. **D)** Relative distance differences between brain mesh and sensors (SQUID MEG minus OPM-MEG). Red colours on the brain mesh indicate closer sensor proximity of OPM-MEG over SQUID MEG. OPM-MEG sensors are visualised as red squares.

### Data collection

Brain activity was recorded using OPM-MEG. OPM-MEG has a closer sensor proximity over cerebellar areas than conventional SQUID-MEG (Figure 1C-D). Participants completed the protocol in a magnetically shielded room as OPM-MEG sensors require the background magnetic field to be below a few nT and are sensitive to surrounding magnetic noise. The OPM-MEG setup comprised 70 sensors (FieldLine HEDscan, FieldLine Inc., Boulder, USA; Figure 1C-D) and data was sampled at 5 kHz. Sixty-eight sensors were allocated to record brain activity and were mounted homogenously in a 3D-printed helmet (with 144 slots available). After helmet preparation, participant’s fiducials and helmet landmark points were digitised using a 3D-registration device (Polhemus FASTRAK, Polhemus Inc., Colchester, USA). We also digitised the participant’s head shape when they did not wear the helmet.

Two of the 70 sensors co-registered the magnetomyography (MMG) of the right trapezius muscle (mounted in slot R117) and magneto-oculography (MOG) of the left eye movements. The MOG and air-puff sensors were attached to Loc-Line plastic arms and located close in front of the left eye. The other ends of the loc-line were mounted in slots L501 and L601 of the helmet respectively. The air-puff nozzle was placed ∼2 cm in front of the eye. To elicit a reflexive eyeblink, the nozzle pressure MPPI-3 pulse injector (Applied Scientific Instrumentation, Eugene, USA) was initially set to 20 psi but was further calibrated if needed. The air puff was triggered through an analogue output generated by DataPixx3 system (VPixx Technologies, Saint-Bruno-de-Montarville, Canada). The DataPixx3 also sent synchronised event triggers to the OPM-MEG data acquisition system, enabling precise co-registration of stimulus timings with the MEG data. Timings of the right index finger responses were measured using a five-button responses box (ResponsePixx, VPixx Technologies, Saint-Bruno-de-Montarville, Canada) and were likewise transmitted to and registered in the OPM-MEG system. Before starting the experimental protocol, resting state brain activity was recorded for five minutes. Participants were instructed to keep their eyes open whilst fixating their gaze. The experiment was concluded by an empty room recording of two minutes. After the experiment, participants completed the experiment awareness questionnaire and Edinburgh handedness questionnaire (Oldfield, 1971).

The experiment awareness questionnaire comprised eleven questions. In five questions, participants were asked for a subjective score of 1-10 on the awareness of task features (1: never noticed the feature; 10: always noticed the feature). Features included awareness on (1) tunnel colour (10 ± 0.75 [median ± interquartile range]), (2) button press (10 ± 0.75), (3) flash (9 ± 1.75), (4) eye blinking during air puff (8 ± 2.75), and (5) eye blinking in absence of air puff (6 ± 1.75; i.e., trials with correct motor responses). Subsequently, they were asked five true/false questions whether (1) the tunnel was circular in shape (false: 14/14), (2) the patterns of the tunnel walls were the same (false: 11/14), (3) the tunnel lightness corresponded to a correct response time (true: 12/14), (4) flash was only presented during some trials (false: 11/14), and (5) T1 and T2 were interleaved at random on each trial (true: 13/14). Finally, they were asked to write down what they believed the response times for the T1 and T2 were (T1: 1.36 ± 0.99 sec [mean ± standard deviation] and T2: 2.50 ± 0.94 sec).

### Data analysis

#### Timings of voluntary movement and conditioned eyeblink responses

All analyses were conducted in MATLAB (version 2022b, MathWorks Inc., Natick, USA). We analysed the timings of the motor and conditioned eyeblink responses (i.e., trials without an air puff administered) in line with Mangili et al. (2025). Both timings were expressed relative to the cue onsets. Note that we refer to the implicit eyeblink response as ‘conditioned eyeblink’ due to its association with the punitive air-puff stimulus as teaching signal, despite differences to the classical eyeblink conditioning paradigm. To identify the conditioned eyeblinks, we specifically focused on the trials in which the participant pressed the button within the omission window. To do so, we used the MOG time series (co-registered as an additional OPM-MEG sensor - see above) as a proxy for eyelid closures and thus to obtain the conditioned eyeblink responses. MOG was low-pass filtered at 10 Hz (second-order bi-directional Butterworth filter) and epoched relative to cue onset from 0 to 1.5 times of the target interval. From the epochs of correct trials (i.e., those with a button press within the omission window), we determined the timing of the conditioned eyeblinks by detecting the minimum of the first time derivative of the MOG. Timings of motor and conditioned eyeblink responses were then analysed over the course of the experiment using a moving average comprising a 30-trial window (15 trials around the trial of interest).

To assess whether we could discriminate the manual and conditioned eyeblink response timings across contexts, we compared the participant-specific averages from the last block (trials 301-450) for both conditions using paired *t*-tests. To test for any learning-related changes across the experiment, for every participant we first obtained the fraction of the correct motor responses and conditioned eyeblinks (binned across 50 trials). To label a trial as having a conditioned eyeblink closure, we considered the standard deviation of the MOG amplitude around the minimum of the first time derivative (time window: -200 to +400 ms). A trial with eyeblink closure was defined as this standard deviation of the MOG amplitude exceeding 2.5 times the standard deviation of the full MOG amplitude epoch (up to 1.2 or 2.4 sec relative to the cue onset). Computing the local standard deviation of the MOG amplitude captures the variability associated with the transient blink response, which can then be compared with the overall variability of the MOG amplitude to determine whether the response is sufficiently pronounced to be classified as a conditioned eyeblink. We then statistically evaluated the response timings between the first (trials 1-50) and last (trials 401-450) bins using paired *t*-tests. We note that the percentages of motor and conditioned eyeblink responses can be correlated as the conditioned eyeblink response was defined on the trials with a correct motor response.

#### OPM-MEG preprocessing

To analyse the OPM-MEG data, we used the open-source FieldTrip toolbox (version 2025-05-23) (Oostenveld et al., 2011). OPM-MEG data were resampled to 500 Hz using an anti-aliasing low-pass filter, demeaned, band-pass filtered between 3 and 200 Hz (second-order bi-directional Butterworth filter) and notch filtered to remove line noise at 50, 100, 150, and 200 Hz. MOG data were high-pass filtered at 1 Hz (second-order bi-directional Butterworth filter) and line noise filtered. We next identified and removed bad channels. Sensors were marked as bad when (I) their standard deviation exceeded three times the mean standard deviation of all channels, (II) their maximum value was greater than 10 times the mean (or median) maximum values across all channels, or (III) their standard deviation was smaller than 100 times the mean standard deviation across all channels. Bad channels were spherically interpolated based on spatially neighbouring channels. We used the positional sensor data as provided by FieldLine Beta2 template model as implemented in FieldTrip. We removed 0 ± [0, 2] (median ± [range]) sensors across the three blocks and all participants. Finally, physiological and non-physiological artefacts were identified and removed using independent component analysis (fastICA algorithm with maximal number of iterations fixed to 5,000). In total, we removed 12.5 ± 8 components (median ± interquartile range across all blocks). Independent components with movement, muscular, and eye artefacts were identified with an automated method. Components were marked as movement or muscular artefactual when their power spectral density revealed a median frequency lower than 1 Hz or higher than 100 Hz respectively. To be labelled as a component with eye artefacts, its topology had to be dominated by frontal sensors. More specifically, this meant that four or more of the eight frontal sensors (L202, R202, L103, R103, L302, R302, L304, and R304) had to dominate the topological weights. Components with muscular, movement, and eye artefacts were verified through visualisation and any additional components with artefacts were added. Finally, we corrected the air-puff timings in the trigger signal by 69 ms. This was the average time delay between the trigger and actual occurrence of the air puff (*n*=98, 69.13 ± 2.51 ms; mean ± standard deviation).

#### OPM-MEG time-frequency analysis at sensor level

To determine how the OPM-MEG activity changes during the trial, we conducted a sensor-by-sensor time-frequency-power analysis. Given that studying cerebellar activity using OPM-MEG has not been studied often (Lin et al., 2019; Lin et al., 2025), we focused on broad time (from -5 to 4 sec relative to the cue onset) and frequency (3 to 80 Hz) ranges. Using single Hanning windows, the time window of each sample was fixed at 3 cycles of the frequency of interest, meaning that the time window length changed as a function of frequency. Time-frequency-power representations were expressed as a relative change by normalising the absolute power to the last second of the inter-trial interval before the context started. Unless stated otherwise, all analyses involving OPM-MEG included trials with both correct and incorrect manual responses.

Participants responded with their right index finger; hence, we focused on the left motor cortex and right cerebellum for the manual responses. The left eye received the air puff and we therefore focus on sensors L115 (over left cerebellum), R115 (right cerebellum), and L308 (left motor cortex). We then determined the statistical significance of the time-frequency-power representations using a surrogate analysis. To do so, we randomly shuffled the samples over time and frequency for 500 surrogates for every sensor. True and surrogate power representations were then compared statistically on a sample-by-sample basis using their analytical distributions. We here employed sample-wise pairwise *t*-tests (FDR with an alpha-level of 0.05) (Maris and Oostenveld, 2007). OPM-MEG activity within time windows was then used for the subsequent source localisation (see below).

#### OPM-MEG source reconstruction

To localise the neural origin of the OPM-MEG activity, we employed Dynamic Imaging of Coherent Sources (DICS) beamformers (Groß et al., 2001). To do so, we first solved the ill-posed inverse problem (i.e., reliably reconstructing the locations of neural sources within the brain from OPM-MEG sensor activity) by estimating a lead-field matrix. Pre-processed OPM-MEG data were then transformed to the frequency domain before obtaining DICS beamformers.

To do so, we computed template volume conduction, source, sensor and lead-field models from the Colin-27 template MRI (Holmes et al., 1998). We did not use individual models because individual anatomical scans were not available (6 out of 14 participants) or fiducial/helmet reference point registrations were inaccurate (4 out of 14 participants). A volume conduction model was estimated as a realistically shaped single shell in line with Nolte (2003). This single shell model was discretised into a source model of 5 x 5 x 5 mm grid. To get the sensor model, we warped the FieldLine Beta2 template model to the fiducial locations of the MRI template. Our sensor model was constrained to all sensors that were consistently available across all participants. Finally, a lead-field matrix was obtained from the volume conduction, source, and sensor models. The lead-field matrix was normalised to avoid centre to the head bias.

We then transformed the OPM-MEG data to the frequency domain. Temporal and spectral parameters were guided by the sensor-level time-frequency representations (Figure 3), where we observed strong beta-band event-related desynchronisation (ERD). We therefore computed DICS beamformers to probe this context-dependent beta-band ERD. Using Slepian sequences, both power and cross-spectral density matrices were estimated for the beta-band ERD (T1 time window: 400 to 900 ms relative to cue onset; T2 time window: 400 to 1,700 ms relative to cue onset). These time windows were selected as these were the largest time windows showing significance at sensor level when contrasting time-frequency power relative to surrogates. We used Slepian sequences with a centre frequency of 21.5 Hz and a spectral smoothing of 8.5 Hz to get beamformers within the beta band (13-30 Hz). For T1 and T2, this resulted in 8 and 21 tapers, respectively. To obtain baseline source power, we estimated power and cross-spectral densities over a time window of 500 (T1) and 1,300 ms (T2) before the end of the inter-trial interval. The length of this time window was matched to the context-specific time window.

DICS beamformers were then estimated from both these inter-trial-interval and context-specific power and cross-spectral densities. This was done on a participant- and context-specific basis. Regularisation of the cross-spectral density matrix was set to 5% (Westner et al., 2022). Filters were constrained to be real valued and have a fixed origin (Westner et al., 2022). Source power at each dipole location were stabilised using a common logarithmic transform, creating log-ratios between source power of the inter-trial interval and context.

To get the context-specific neural sources, we created a statistical contrast between the voxel-by-voxel source power of the inter-trial interval and context (paired *t*-test with an false-discovery-rate-controlled [FDR] alpha-levels of 0.001) (Maris and Oostenveld, 2007). The derived statistical masks were used to visualise the context-specific power differences. Statistical (*t*-values) and source-power volumes were visualised using the MRIcroGL software (https://www.nitrc.org/projects/mricrogl) (Rorden, 2025).

#### Source-reconstructed time series

As it is not trivial to relate sensor-level signals to brain anatomy, we estimated source-space time series by creating so-called virtual channels using DICS-beamformers that were estimated over broad frequency (3-80 Hz) and time (-5 to 4 s relative to cue onset) ranges. These source-reconstructed time series at regions of interest (ROIs) can then be used for our hypothesis testing. We determined these ROIs from the Automated Anatomical Labelling (AAL) atlas (Tzourio-Mazoyer et al., 2002). Given that we expected task-relevant responses in the left cortical motor areas and bilateral cerebellar areas, we selected the left paracentral lobule and all cerebellar areas as our ROIs. From the AAL atlas, we evaluated which bilateral cerebellar region showed the strongest beta-band ERD, which was the combined left and right cerebellar lobule VI (Table S1). We note that taking the mean beta-band power rather than the maximal power also led to selecting bilateral cerebellar lobule VI (Table S1). To reconstruct source-level activity at cerebellar lobule VI as well as left precentral lobule (left motor cortex), we first estimated broad-band filters using beamformers using similar parameters as taken for the beta-band beamformers. Participant-specific filters were estimated with all data from both contexts. The latter is to avoid that spatial filters will be different between the contexts. Sensor-level data were then filtered using these broad-band filters at voxels that exhibited maximal beta-band ERD (as determined from the statistical contrasts). Source-level data of bilateral cerebellar lobules VI and left precentral lobule were used in subsequent analysis.

#### Source-reconstructed time-frequency analysis and statistics

Using the source-reconstructed activity of bilateral cerebellar lobule VI and left precentral lobule, we then examined the time-frequency power at these areas after the cue onset. Again, we created on broad-band representations in time (from -5 to 4 sec relative to the cue onset) and frequency (3 to 80 Hz) ranges. Hanning window lengths were set to 3 cycles of the frequency of interest. Resulting time-frequency-power representations were normalised to the last second of the inter-trial interval.

We then determined the statistical significance of the time-frequency-power representations using a surrogate analysis. Similar to sensor level, we randomly shuffled the samples over time and frequency for 500 surrogates for every time-frequency representation. True and surrogate power representations were then compared statistically on a sample-by-sample basis using their analytical distributions (sample-wise pairwise *t*-tests; FDR with an alpha-level of 0.05) (Maris and Oostenveld, 2007). Sample-by-sample paired *t*-tests were also employed to compare time-frequency power across ROIs and across contexts. Statistical contrasts across ROIs were left precentral lobule against bilateral cerebellar lobules VI (done separately for both cerebellar hemispheres). Again, alpha levels were set to 0.05 (FDR corrected).

#### Cortical and cerebellar learning dynamics

We next tested the hypothesis that cerebellar learning precedes cortical learning. To do so, we statistically evaluated whether cerebellar and cortical beta-band ERD changed differently throughout the course of the experiment. First, the time-frequency-power representations of left precentral lobule and bilateral cerebellar lobules VI were averaged over frequencies (13 and 30 Hz) and time (300-2,000 ms after the cue onset). This time window was guided by the statistical comparisons against surrogates. Time-frequency-power representations were additionally masked by this statistical contrast. Trial-by-trial beta-band dynamics were then pooled into bins of 30 trials irrespective of the context conditions. We then constructed a linear mixed-effects model to statistically evaluate power differences in training rates for the left precentral lobule and bilateral cerebellar lobules VI. In this model, the fixed effects *Bin* (main), *ROI* (main), and *Bin*-by-*ROI* (two-way interaction) were added as well as a random effect of *Participant*, which acted on both intercept and slopes (*Bin* and *ROI*). Alpha-level was set at 0.05.

#### Correlative analysis between cerebellar power timings and motor timings

To test how the cerebellar beta-band ERD related to our behavioural readouts, we studied associations between cerebellar beta-band timings and manual/eyeblink timings. To do so, we constructed four linear-mixed effects models. Beta-band timing of either left or right cerebellar lobule VI was included as a fixed effect. Either manual or conditioned eyeblink timings was included as the response variable. *Participant* was included as random effect, acting on both intercept and slope of the beta-band timing. We also report the partial correlation coefficient to provide an index for the effect size. Beta-band timing was defined as the timing when 50% of the area under curve was reached (time window between 300 and 2,000 ms). This timing was then expressed relative to the correct timings at 800 and 1,600 ms for T1 and T2, respectively (where negative values correspond to a beta-band timing that occurred earlier than the manual and conditioned eyeblink timing). Manual and eyeblink timings were similarly normalised to the correct timings at 800 and 1,600 ms. Since cerebellar power was averaged over 30 trials, we did the same for the manual and eyeblink timings.

#### Estimating cortico-cerebellar coupling through partial directed coherence

Finally, we examined context-specific functional connectivity between the source-reconstructed activity of the left precentral lobule and bilateral cerebellar lobule VI. To do so, we employed partial directed coherence which is a directional coupling metric (Baccalá and Sameshima, 2001). Briefly, for partial directed coherence, we first estimated the multivariate autoregressive coefficients with model order 8 (equalling 16 ms given the sampling frequency of 500 Hz). This is In line with time conduction delays of cortico-cerebellar loops (Bloedel and Courville, 1981). Coefficients were derived within time windows of 1,000 ms around the sample of interest (-500 to +500 ms). Next, the transfer function, innovation noise covariance and cross spectrum matrices were derived from the autoregressive coefficients, essentially Fourier transforming the autoregressive coefficients. From these Fourier-transformed components, we then estimated partial directed coherence as proposed by Baccalá and Sameshima (2001). To test the hypothesis for statistically significance coupling, we first constructed surrogate coupling. To do so, for each time sample we randomised the Fourier phases after transforming the 1,000-ms epoched data to the frequency domain (Faes et al., 2009). Data were then transformed to the time domain using the inverse of the Fourier transform (Faes et al., 2009). This surrogate approach retains the original signal properties whilst destroying the coupling through phase randomisation (Faes et al., 2009). Surrogate coupling was estimated for each surrogate (100 surrogates in total). True and surrogate coupling representations were then compared statistically on a sample-by-sample basis using their analytical distributions. We here employed sample-wise pairwise *t*-tests (FDR with an alpha-level of 0.05).

## Results

### Manual and eyeblink responses exhibit context-specific timings

To investigate context-dependent manual and eyeblink timings in this behavioural paradigm that yoked incorrectly timed right-hand button presses to left-eye air puffs, we initially replicated the response timings established by Mangili et al. (2025). We found that the manual response and eyeblink times were closely matched to the expected ones at 800 (T1) and 1,600 (T2) ms (Figure 2A-D; Figures S1-S4). For the last block (consisting of trials 301-450), the contextual manual response times were significantly different between contexts (*t*(13): 3.89, *p*: 0.002; Figure 2E). For both contexts, the percentage of correct manual responses also increased over the course of the experiment (*t*(13): 3.40, *p*: 0.005; Figure 2F). We then assessed the conditioned eyeblink responses. Focusing on the correct manual trials when no air puff was presented, we found that the timing of the conditioned eyeblink response was also context specific (*t*(10): 14.57, *p*: 5e-8; Figure 2G). The percentage of conditioned eyeblink responses increased over the course of the experiment (*t*(13): 3.42, *p*: 0.005; Figure 2H; note that due to our definition the percentage of conditioned eyeblink responses can never be higher than the percentage of correct manual responses). Collectively, this means that the manual and conditioned eyeblink responses matched the timings of T1 and T2.

**Figure 2.**
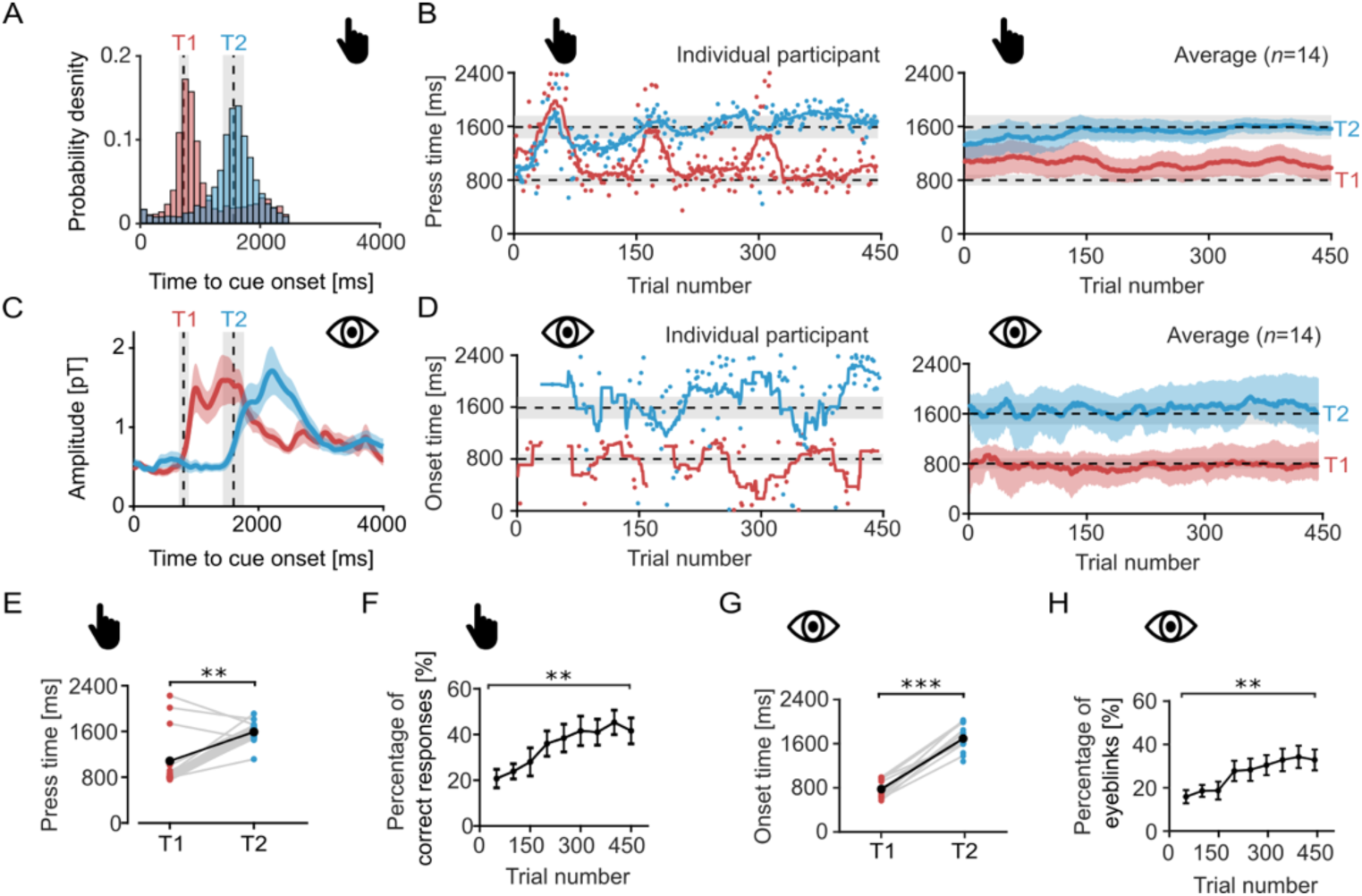
Task design and behavioural findings. **A)** Probability densities of manual responses of all trials and participants. **B)** Manual response timings (relative to cue onset) when the button was pressed for T1 and T2. Timing-over-trial graphs are moving averages (+/-15 trials) for one participant (top panel) and all participants (n=14; mean and standard error of the mean). Grey patches are the omission windows for both contexts. **C)** Grand average magneto-oculographic responses for T1 and T2. Responses are visualised as means and standard errors of the mean (shaded surfaces). Vertical grey patches represent the respective omission windows. **D)** The conditioned eyeblink onset of the left eye (similarly structured as panel B). **E)** Inter-context comparisons of the mean manual response timings across the last block (trials 301-450). Red (T1) and blue (T2) samples are individual participants, and black samples are group-specific means for both contexts. **F)** Percentages of correct manual responses evaluated over the course of the experiment. Black dots and error bars are mean and standard error of the mean across participants. The statistical evaluation is the percentages of correct responses between trials 1-50 and 401-450. **G-H)** Panels for the conditioned eyeblink timings and percentage of eyeblinks (which are similarly structured as panels E and F, respectively). **: p: <0.01; ***: p: <0.001.

### Beta-band ERD exhibits context-dependent timing across the contralateral motor cortex and bilateral cerebellum

We examined OPM-based oscillatory dynamics in sensors over left (contralateral) cortical motor and bilateral cerebellar regions. Bilateral cerebellar regions were examined as these areas were expected to be involved in the manual response (right hemisphere) and the eyeblink (left hemisphere), respectively. First, sensor-level time-frequency-power representations revealed context-dependent beta-band ERD over the left cortical motor sensors and bilateral cerebellar sensors (Figure 3A and C; Figures S5-S6). Beta-band ERD occurred following the flash and temporally aligned with the omission window. Specifically, beta-band ERD at the cortical motor sensor was observed at narrow windows before the respective omission windows (T1: 556-656 ms; T2: 1,144-1,420; *p*: <0.05 FDR corrected; sample-wise dependent *t*-tests relative to 500 surrogates). Beta-band ERD at sensors over the left and right cerebellum was also observed at longer time windows (left cerebellar sensor: 328-1,122 ms [T1] and 296-1,780 ms [T2]; right cerebellar sensor: 316-1,122 ms [T1] and 294-1,740 ms [T2]; *p*: <0.05 FDR corrected; sample-wise dependent *t*-tests relative to 500 surrogates). Context-dependent beta-band ERD was widespread, with maximal effects over posterior sensors (Figure 3B and D; Figures S7-S8). Neither effect was attributable to eyeblinks, neck muscle activity (Figure S9) or air puffs delivered to the eye in incorrect trials, as restricting the analysis to correct trials only, still yielded robust beta-band desynchronisation across sensors over the left primary motor cortex and bilateral cerebellar regions (Figure S10). In sum, we found that sensors over the left cortical motor and bilateral cerebellar areas displayed context-dependent beta-band ERD, which temporally scaled to T1 and T2.

**Figure 3.**
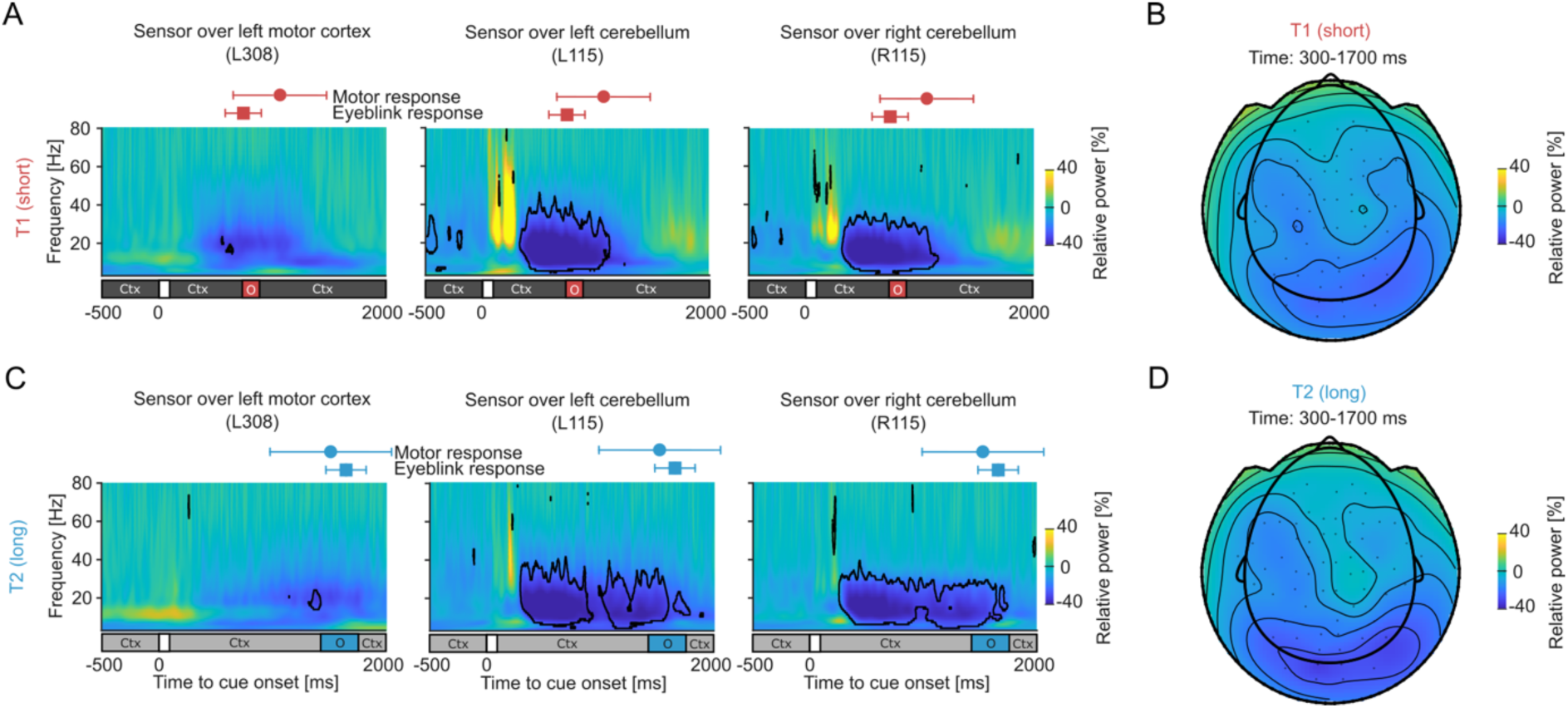
Time-frequency-power representations and topographic plots. **A)** Time-frequency-power representations of the T1 for sensor L308 (left motor cortex), L115 (left cerebellum) and R115 (right cerebellum). Coloured circles and squares are the mean times of the manual and conditioned eyeblink response events (error bars are standard deviations over participants). Bar below the time-frequency representation indicates the timings within the trial. The context was followed by a brief flash of 100 ms after which the tunnel started moving. Black outlines indicate statistical significance (p: <0.05; false discovery rate). Ctx: context; O: omission window. **B)** Topological map of the beta-band desynchronisation of T1 after the cue onset (time window: 300-1,700 ms). **C-D)** Time-frequency results for T2 (panels are similarly structured as panels A and B).

Posterior sensors also exhibited post-cue gamma-band (30-80 Hz) synchronisation. However, source localisation placed these signals outside anterior cerebellar regions related to manual and eyeblink regions (lobules V-VI), instead projecting to the visual cortex (Figure S11). Furthermore, gamma-band synchronisation was more pronounced after a cue flash following a dark tunnel (T1) relative to a light tunnel (T2) and also occurred at trial onset following static tunnel presentation in the absence of manual responses (Figure S12). Together, these findings indicate that post-cue gamma dynamics predominantly reflect sensory visual processing.

In addition to the hypothesised beta-power modulations, we observed alpha-band desynchronisation over comparable time windows in posterior sensors. Given the established role of alpha rhythms in visual processing, we examined whether these observed responses reflected spatial smearing or spectral leakage from the visual cortex into cerebellar reconstructions. For our initial time-frequency analysis, the Hanning window length was fixed at 3 cycles per frequency of interest. To test whether increasing window length would improve frequency resolution and separate alpha-from beta-band desynchronisation, we re-analysed the data using 7-cycle windows. Time-frequency power remained characterised by statistically significant, co-occurring alpha- and beta-band desynchronisation (*p* < 0.05, FDR-corrected; Figure S13), demonstrating that beta-band desynchronisation does not merely reflect spectral leakage from lower alpha frequencies. Furthermore, source-level beamforming in the alpha band localised power changes between the context window and the inter-trial interval predominantly to cerebellar regions rather than visual cortex (Figure S14). Together, these control analyses confirm that posterior alpha-band desynchronisation reflects true cerebellar activity rather than spatial smearing or visual processing artifacts induced by the visual flow stimulus.

Consistent with right-hand button press and implicit left eyeblink responses, the context-dependent beta-band ERD was source-localised to left (contralateral) cortical motor and bilateral cerebellar areas for T1 and T2, respectively (*p*: <0.001 FDR corrected; Figure 4; sample-wise dependent *t*-tests relative to inter-trial interval). When creating a common source localisation for both contexts (used for the construction of virtual channels), within the cerebellum the strongest beta-band ERD was localised to the bilateral lobules VI (Table S1). ERD was found in contralateral cortical motor areas (Brodmann areas 4 and 6), however the extent and magnitude of the ERD in the motor cortex was less pronounced than expected.

**Figure 4.**
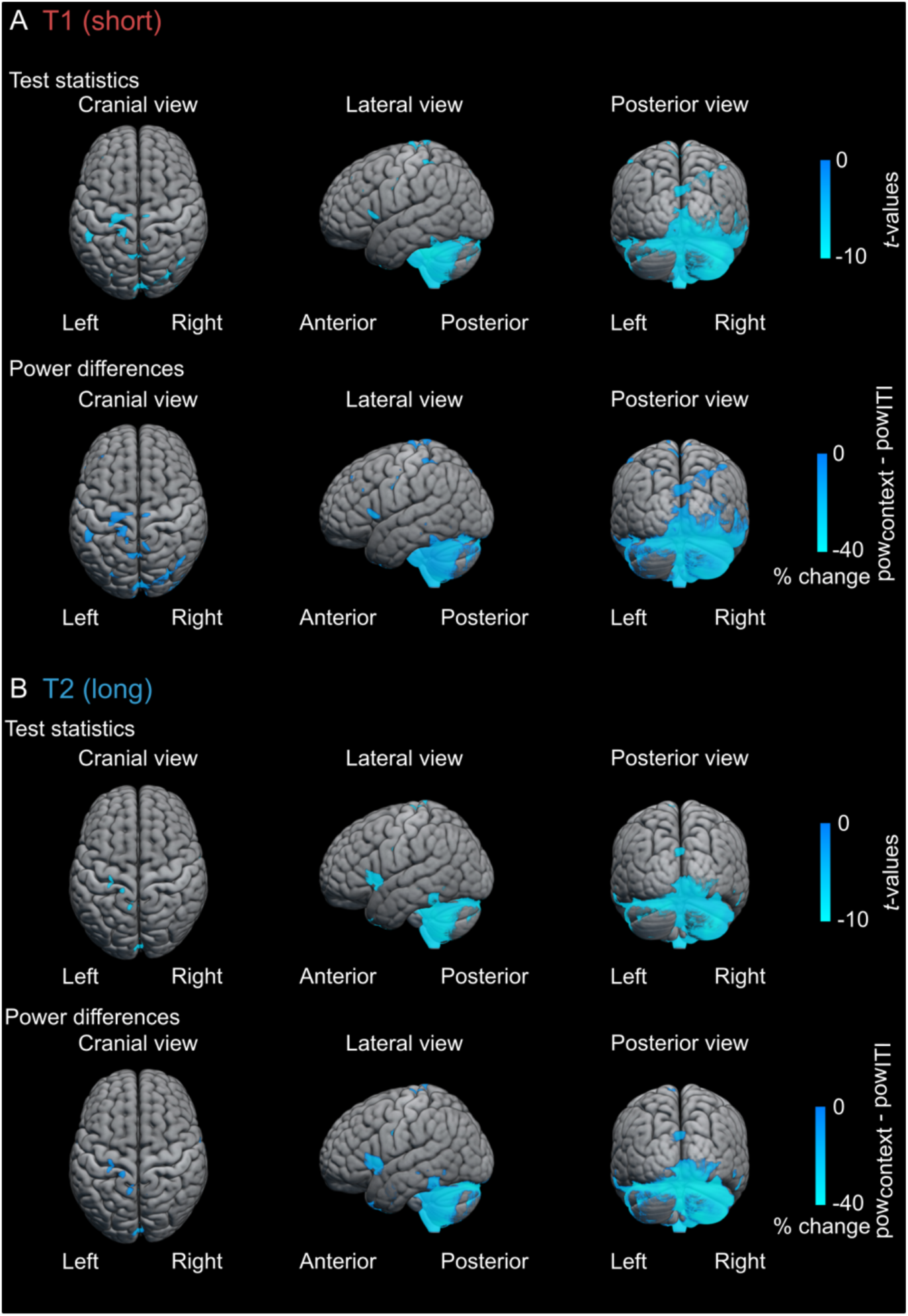
Spatial localisations of A) T1 and B) T2. Source localisations are constrained to beta-band frequencies (frequency range: 13-30 Hz) and context (time window T1: 400 to 900 ms; time window T2: 400 to 1,700 ms). These time windows were derived from the sensor-level time-frequency statistics. Test statistics and source-power differences are masked by the statistical contrasts between context and inter-trial interval (p: <0.001 false discovery rate).

### Training-related changes in cerebellar desynchronisation precede cortical changes

In line with sensor-level results, source-reconstructed beta-band ERD in the left motor cortex and bilateral cerebellar lobules VI occurred after each cue and temporally aligned with the context-guided target press intervals. Significant time windows (*p*: <0.05 false discovery rate [FDR] corrected; sample-wise dependent *t*-tests relative to 500 surrogates) were 400 to 1,256 ms (T1) and 742 to 1,720 ms (T2) over the left precentral lobule (Figure 5A). Beta-band ERD at the left and right cerebellar sensors was observed at longer time windows (Figure 5A; left cerebellar lobule VI: 334 to 1,214 ms [T1] and 298 to 1,836 ms [T2]; right cerebellar lobule VI: 318 to 1,330 ms [T1] and 300 to 1,932 ms [T2]; *p*: <0.05 FDR corrected; sample-wise dependent *t*-tests relative to 500 surrogates) and was more pronounced than in the motor cortex (Figure S15; *p*: <0.05 FDR corrected; sample-wise dependent *t*-tests). We note that we find comparable time-frequency power when taking the mean spatial filters of the ROIs rather than the maximal filters (Figure S16).

**Figure 5.**
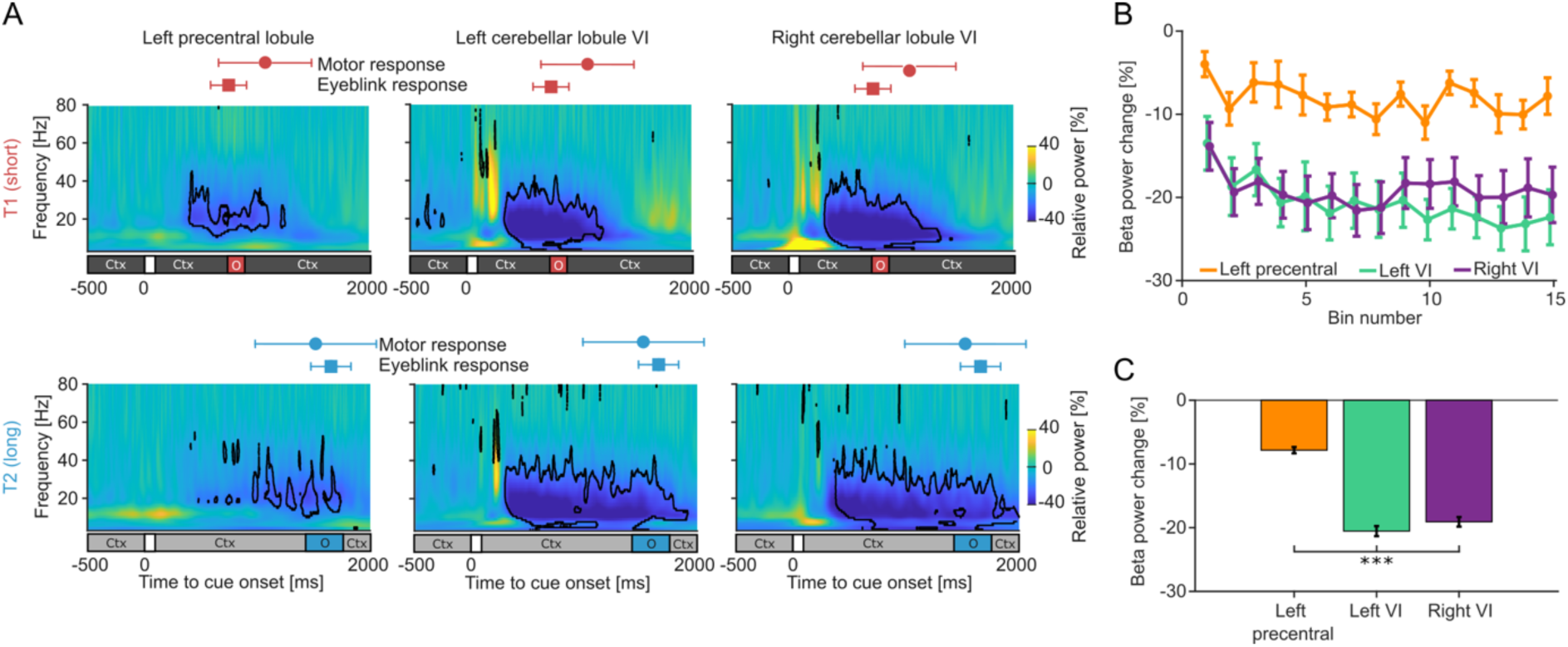
Source-reconstructed time-frequency power and training-related beta-band power changes. **A)** Time-frequency-power representations of T1 and T2 for the left paracentral lobule and bilateral cerebellar lobules VI. Coloured circles and squares are the mean times of the manual and conditioned eyeblink response events (error bars are standard deviations over participants). Black outlines indicate statistical significance (p: <0.05; false discovery rate). Bar below the time-frequency representation indicates the timings within the trial. The static context was followed by a brief flash of 100 ms after which the context started moving. Ctx: context; O: omission window. **B)** Training-related changes in beta-band desynchronisation for the regions of interest (ROIs). Changes are expressed relative to the inter-trial interval and pooled into 30-trial bins. Bars and error bars represent group means and standard errors of the mean over participants. **C)** Main effect of ROI. Bars and error bars represent regions means and standard errors of the mean over bins and participants. ***: p: <0.001.

Behavioural evidence from implicit eyeblink timing indicates that the cerebellum might be capable of rapid, flexible motor learning (Mangili et al., 2025). Hence, we tested whether training-related changes in beta-band ERD differed across cerebellum and motor cortex. To test this hypothesis, trial-by-trial beta-band power from the left precentral lobule and the bilateral cerebellar lobules VI were pooled into 30-trial bins. This revealed a significant *bin*-by-*ROI* interaction (F(2,600): 5.21, *p*: 0.006; Figure 5B) which generalised across all tested bin sizes (also see Figure S17). The motor cortex showed relatively stable beta-band ERD across bins, whereas both cerebellar regions exhibited stronger reductions over time (Figure 5B). For the model comprising 30-trial bins, we also found a main effect of *ROI* (F(2,600): 9.80, *p*: 6e-5; Figure 5C), suggesting that beta-band ERD is more pronounced relative to the inter-trial interval in cerebellar compared to cortical motor areas. Taken together, these results indicate that beta-band ERD becomes more pronounced in the cerebellum with training.

### Cerebellar beta-band ERD timing is predictive of manual response timing

To determine how cerebellar beta-band ERD relates to motor execution, we evaluated its association with manual and eyeblink response timing (Figure 6 and Figure S18). We hypothesised that bilateral lobules VI play a different, hemisphere-specific role. That is, the ipsilateral (right) lobule should be related to the manual response only, whereas the contralateral (left) lobule which processes the periocular air puff that delivered to the left eye should relate to both the conditioned left eyeblink response and the right-hand manual response, because the latter is timed to avoid the air puff. As expected, beta-band timing (defined as the timing when 50% of the area under curve relative to T1/T2) in the ipsilateral (right) lobule VI was correlated with the manual response timing (*t*(200): 4.51; *p*: 1e-5; *r*: 0.30; Figure 6B), but not with the conditioned eyeblink timing (*t*(192): 0.97; *p*: 0.34 *r*: 0.07; Figure 6E). However, while the beta-band timing in the left cerebellar lobule VI was correlated with the manual response timing, possibly reflecting the yoked task (*t*(200): 4.84; *p*: 3e-6; *r*: 0.32; Figure 6C), it was not related to the conditioned eyeblink response timing (*t*(192): 0.71; *p*: 0.48; *r*: 0.05; Figure 6D). These positive correlations between beta-band and manual timings were qualitatively replicated for different bin sizes of 10, 15, 25, 50, 75, and 150 trials. These results suggest that bilateral cerebellar activity is primarily involved in the timing of the manual responses which determine whether an air puff is delivered in this paradigm; however, we did not find a clear association of the left lobule VI with the implicit eyeblink response.

**Figure 6.**
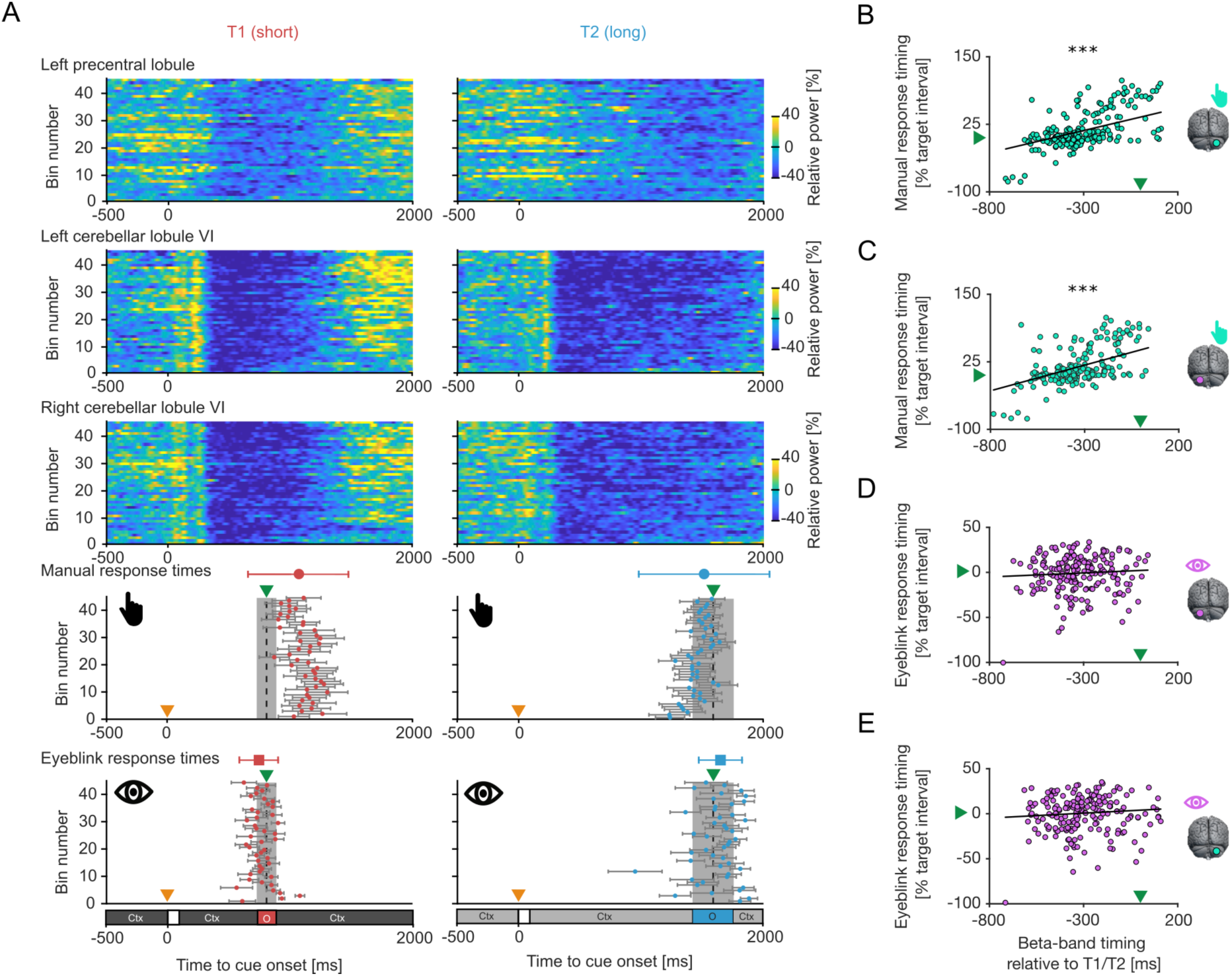
Associations between timings of power-related and behavioural outcomes. **A)** Time-resolved power changes for the left precentral lobule and bilateral cerebellar lobules VI for both contexts as well as the timings of the manual and conditioned eyeblink responses. Power is averaged over the frequencies of 13 and 30 Hz (comprising beta-band frequencies) and binned over every 5 trials. Manual response and conditioned eyeblink times are the mean and standard error of the mean. Bar below the conditioned eyeblink response times indicates the timings within the trial. Orange and green triangles indicate the timings of the cue onset and T1/T2, respectively. The static context was followed by a brief flash of 100 ms after which the context started moving. Ctx: context; O: omission window. **B-E)** Linear mixed-effects models between beta-band timing of the bilateral cerebellar lobules and timings of the **B-C)** manual and **D-E)** conditioned eyeblink responses. Every point comprises an average of 30 trials. The black graph is the linear fit for the mixed-effects model. Green triangle is analogous to panel A and indicate the time interval. ***: p: <0.001.

### Cortico-cerebellar coupling is bidirectional at beta-band frequencies

Finally, we tested whether cerebellar activity and cortical motor activity are coupled to enable yoked learning. To do so, we first examined how the beta-band power covaried across regions of interest. We found that the beta-band power in the contralateral motor cortex activity was correlated with that of the bilateral cerebellar activity (right: *t*(200): 2.47; *p*: 0.01; *r*: 0.17; Figure 7A and left: *t*(200): 2.11; *p*: 0.04; *r*: 0.15; Figure 7B), demonstrating that the beta-band changes covary across both regions.

**Figure 7.**
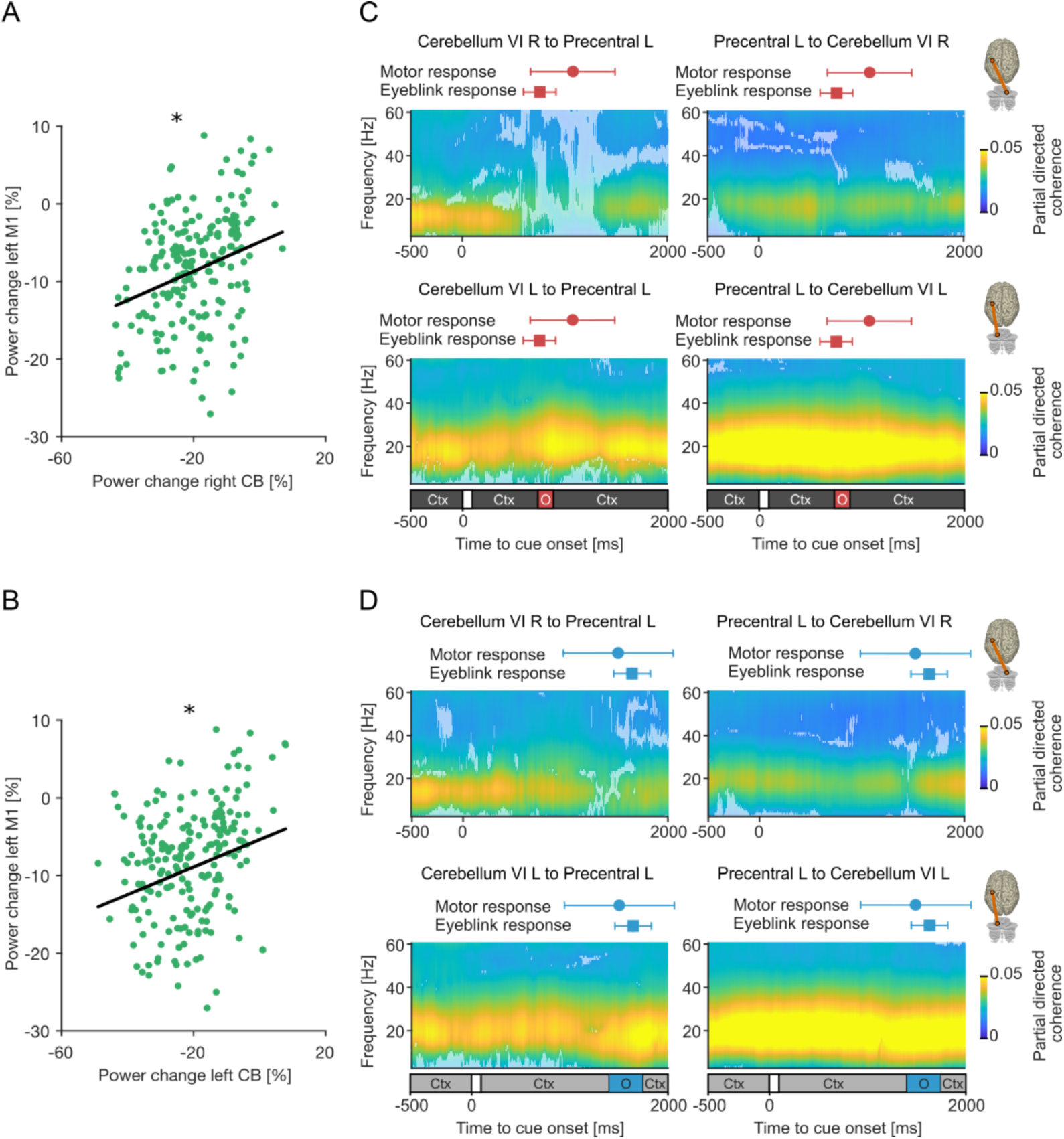
Bidirectional cortico-cerebellar coupling. Linear mixed-effects models between power changes in bilateral cerebellum (CB) and left cortical motor (M1) areas **A)** right cerebellum; **B)** left cerebellum). Every point comprises an average of 30 trials. The black graph is the linear fit for the mixed-effects model. Partial directed coherence between the left paracentral lobule and bilateral cerebellar lobules VI for **C)** T1 and **D)** T2. Opaque samples indicate statistical significance relative to surrogate coupling (p: <0.05; false discovery rate, sample-wise dependent t-tests relative to 100 surrogates). Coloured circles and squares are the mean times of the manual and conditioned eyeblink response events (error bars are standard deviations over participants). Bar below the time-frequency representation indicates the timings within the trial. The static context was followed by a brief flash of 100 ms after which the context started moving. Ctx: context; O: omission window. *: p: <0.05.

To explore the temporal dynamics of coupling throughout the trial, we further examined the cortico-cerebellar coupling by employing partial directed coherence. This directed coupling metric revealed cortico-cerebellar coupling at beta-band frequencies. Coupling was bidirectional from cerebellum to cortex and *vice versa* (*p*: <0.05, FDR corrected; sample-wise dependent *t*-tests relative to 100 surrogates; Figure 7C-D). Directional coupling between ipsilateral (right) cerebellar lobule VI and contralateral (left) motor cortex, both anatomically linked to the manual response decreased specifically at the time of the omission window. This was not found for the directional coupling from contralateral (left) motor cortex to ipsilateral (right) cerebellar lobule VI. No decreases in coupling were found between the left cerebellar lobule VI and left motor cortex, or *vice versa* (also see Figure S19). It is unlikely that this decrease in coupling in the cortico-cerebellar regions related to the manual response can solely be explained by power decreases, since beta-band ERD was observed for both cerebellar lobules at comparable time windows (Figure 5).

A direct comparison of coupling directionality (i.e., cortex to cerebellum, or cerebellum or cortex) did not yield an effect (*p*: >0.05, FDR corrected; Figure S20A). We also did not find statistical differences across contexts (*p*: >0.05, FDR corrected; Figure S20B) and cerebellar regions (i.e., coupling for right and left cerebellum with precentral area; *p*: >0.05, FDR corrected; Figure S20C).

## Discussion

Accurate and flexible movement timing is a cornerstone of adaptive behaviour. However, how distributed cortico-subcortical networks, particularly the cortico-cerebellar loops, interact to achieve this skilled temporal control remains a central question in motor neuroscience (Hazeltine et al., 1997; Nobre et al., 2007; Kornysheva, 2016; Remington et al., 2018; Stine and Jazayeri, 2025). To record cerebellar and cortical activity simultaneously, we implemented high-density OPM-MEG during a contextual learning task that yoked incorrectly timed right-hand button presses to left-eye air puffs (Mangili et al., 2025). Across both the contralateral motor cortex and bilateral motor cerebellar lobules, we observed context-dependent reductions in beta-band ERD that temporally scaled to T1 and T2. Their timing predicted the manual response and became more pronounced with training in the cerebellum than motor cortex. Importantly, contralateral cortical and bilateral cerebellar activity showed beta-band-specific bidirectional coupling, whereas movement execution was characterised by a transient weakening of coupling from the hand-related ipsilateral cerebellum to the contralateral motor cortex around the time of the manual response.

### Cerebellar beta-band ERD as a marker of inhibition-excitation dynamics

Beta-band ERD is a hallmark of motor control processes (Pfurtscheller and Da Silva, 1999). According to the “status-quo” hypothesis, beta-band oscillations are expressed more strongly during maintenance of the current (sensorimotor) state and are attenuated when a change in motor state is required (Engel and Fries, 2010). Specifically, beta-band oscillations in the motor cortex and basal ganglia are associated with GABAergic inhibition (Groth et al., 2021), whereas beta-band ERD serves to permit the disinhibition and subsequent release of a motor plan (Barone and Rossiter, 2021). We show that in addition to beta-band ERD in the motor cortex (contralateral to the manual response), a pronounced motor-related beta-band ERD can be found in the cerebellum, appearing strongest in the bilateral lobules VI. These right and left cerebellar lobules are anatomically linked to the hand performing the manual response and the eye receiving the air puff respectively. Whilst this beta-band ERD was prominently observed in the left motor cortex and bilateral cerebellum, it was unexpectedly stronger in cerebellar compared to cortical areas. Sensor coverage over primary motor and premotor regions was restricted in our OPM setup relative to conventional SQUID-MEG recording configurations, which may have reduced sensitivity to the full magnitude of the cortical response.

At the microcircuit level, this macroscopic beta-band ERD may reflect the disinhibition of the deep cerebellar nuclei (DCN) by upstream inhibitory Purkinje cells. These nuclei are disinhibited via a transient, coordinated reduction in Purkinje cell firing, which is a well-documented mechanism in classical eyeblink conditioning (Medina et al., 2000; Ten Brinke et al., 2015; Koppen et al., 2026). Since Purkinje cells are geometrically aligned parallel to one another and perpendicular to the cerebellar cortex, their architectural orientation relative to the scalp mimics that of pyramidal cells in the cerebral cortex, making their population-level open-field dipoles highly detectable via surface OPM-MEG arrays (Andersen et al., 2020; Roos et al., 2026).

The intra-trial power dynamics revealed a cerebellar involvement in motor preparation in addition to execution. Trial-by-trial cerebellar beta-band ERD started earlier and was more prolonged than cortical beta-band ERD. These temporal profiles argue against cerebellar computations associated with motor execution only. Instead, they point to an active feedforward mechanism operating during motor preparation and accurately release the timed motor command according to a particular sensorimotor context (Gao et al., 2018; Sinha et al., 2025b). At the microcircuit level, Purkinje cell firing profiles within cerebellar lobule IV/V have been shown to directly reflect the prior temporal probability of air-puff stimuli in an eyeblink conditioning task (Koppen et al., 2026). In contrast, beta-band ERD in the motor cortex reflects the disinhibition of layer V pyramidal populations that increase their firing rate to drive the manual response, operating on a more transient timescale tightly bound to movement execution (Figure 8A) (Barone and Rossiter, 2021).

**Figure 8.**
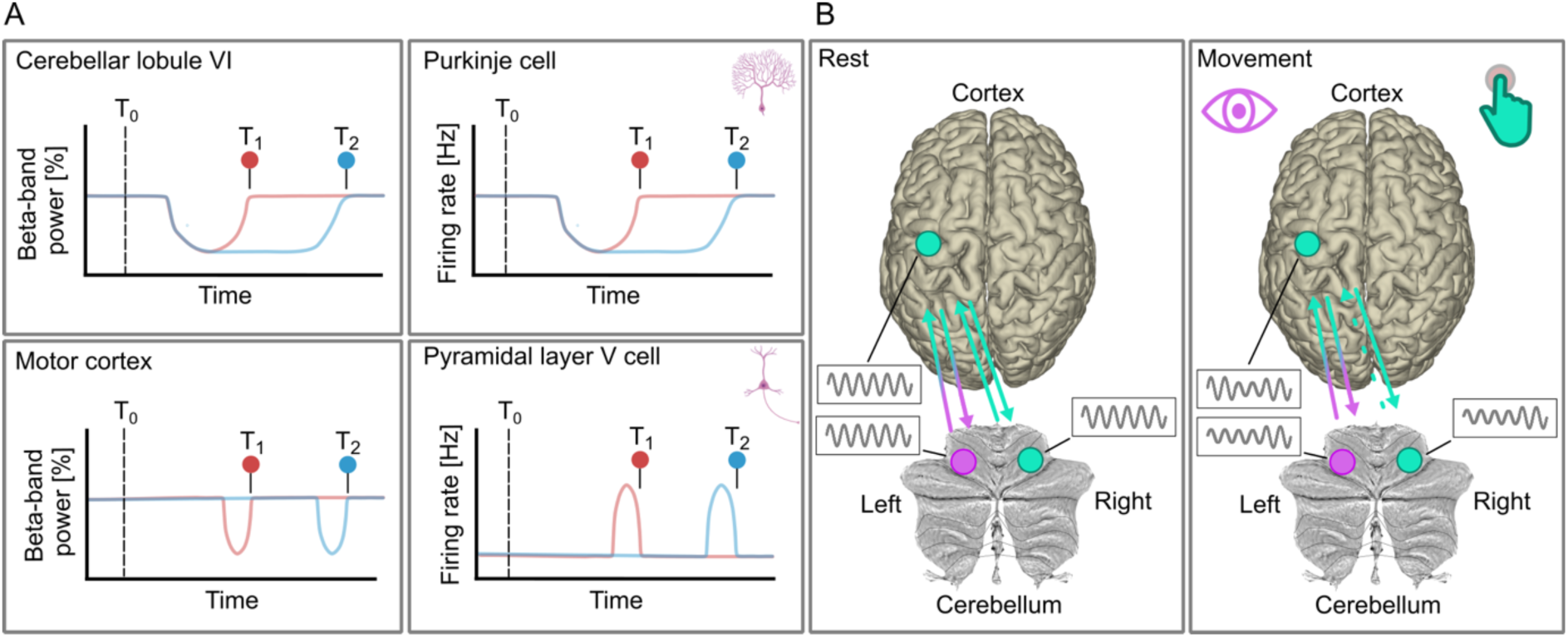
Schematic model of microcircuit and macroscopic correlates of cortico-cerebellar dynamics during context-dependent motor timing. **A)** Proposed mapping between macroscopic OPM-MEG signatures (left panels) and cellular circuits (right panels) across two context-dependent target intervals (T_1_ and T_2_). Anticipating T_1_ and T_2_, OPM-based beta-band activity decreases in the cerebellar layer VI relative to a cue provided at t_0_. It is thought that this reflects firing rate reduction of Purkinje cells at the cellular level whose firing rate reduction is temporally scaled to the target interval. Conversely, macroscopic cortical motor beta-band power and cortical layer V pyramidal cells exhibit a desynchronisation restricted to a window closer to the target timings at T_1_ and T_2_. **B)** Schematic of changes in cortico-cerebellar coupling. Left and right cerebellum are involved in the control of the left eye (in purple) and right hand (in cyan). The left motor cortex interacts with bilateral cerebellar networks during resting state, as shown through bidirectional coupling at beta-band frequencies. The cyan line from right cerebellum to left motor cortex denotes the classical manual motor pathway tracking execution, displaying a weakening of coupling around movement execution. The flat cerebellar surface was obtained from the SUIT Toolbox (Diedrichsen and Zotow, 2015).

In addition to the hypothesised beta-band modulations, we unexpectedly observed alpha-band desynchronisation in posterior sensors. Because visual flow characteristically suppresses visual cortical alpha rhythms (Bauer et al., 2014), we tested whether this response reflected spatial smearing or spectral leakage from visual cortex. Re-analysing the data with longer 7-cycle time windows confirmed that alpha- and beta-band desynchronisations remained distinct and co-occurring (Figure S13). Furthermore, source-level beamforming localised alpha-band power changes predominantly to cerebellar lobules rather than visual cortex (Figure S14). These findings rule out visual-flow-induced cortical artifacts, confirming that posterior alpha-band modulations reflect intrinsic cerebellar activity alongside the motor-related beta response, suggesting that alpha-band modulations reflect a co-occurring cerebellar process that operates alongside or complementary to motor-related beta ERD during temporal control.

### Cerebellar-cortical coupling as a gating mechanism

In the current paradigm, air-puff stimulation served as an aversive feedback signal triggered by incorrect manual response timing. This design rendered contextual implicit cerebellar timing computations directly relevant to the manual motor output (Mangili et al., 2025). Crucially, while the air puff also evoked an eyelid closure, this conditioned response had no causal bearing on the probability of the delivery of the air-puff stimulus itself. Our data provide first direct neural evidence for this behavioural yoking between an implicit cerebellar-mediated motor timing and an explicit cortically mediated manual timing, demonstrating that bilateral, not just manual response-related ipsilateral, cerebellar beta-band ERD correlates robustly with trial-by-trial manual motor timing accuracy. In our data, left cerebellar beta-band ERD anatomically mapped to left eyeblink responses correlated directly with left cortical motor beta-band ERD, which drives the right-hand manual response, suggesting potential cross-effector relevance of the ERD in an eyeblink-related cerebellar region. However, the precise pathways that relay this predictive timing signal from the left cerebellum to the right cerebellar hemisphere and/or the left motor cortex remain to be determined. It is plausible that this cross-effector yoking of the air-puff signal to the manual response system is enabled progressively across trials and via a third region which we did not identify in the OPM-MEG recordings.

Importantly, our findings reveal the dynamics of the trial-by-trial cortico-cerebellar communication in the beta band during timed motor execution. While baseline connectivity between bilateral cerebellar lobules VI and left motor cortex was characterised by stable, bidirectional beta-band coupling, this state shifted temporarily during movement-relevant time windows. Specifically, preparing a manual movement triggered a directed decrease in beta-band coherence from the right cerebellum to the contralateral left motor cortex (Figure 8B), the canonical circuit controlling the right-hand response. This transient, unidirectional drop in beta coupling may reflect a gating mechanism: by breaking baseline tonic synchrony to retain the ‘status quo’ (Engel and Fries, 2010), the circuit releases motor cortex from a holding state, enabling disinhibited cerebellar output to drive movement execution. This coherence drop is unlikely to be a result of local power loss, as beta power in the left eyeblink-related cerebellum also decreased without altering coupling. Although caution is needed in interpreting these partial directed coherence findings as they cannot confirm true causality of the neural coupling, they are in line with recent non-human primate studies. These studies show that blocking the output of the DCN to the cortex reduces firing in the motor cortex causes beta-band ERD in the motor cortex to be less pronounced (Israely et al., 2025; Sinha et al., 2025b) and muscle activity and effector velocity during reaching tasks to be reduced (Sinha et al., 2025a).

This transient reduction in directed beta-band coupling provides a striking parallel to invasive electrophysiological recordings in human patient populations examining cortico-basal-ganglia coupling. Recordings from scalp-based MEG/EEG and deeper-brain local field potentials (LFP) in the subthalamic nuclei and globus pallidus demonstrate that continuous beta-band coupling between the motor cortex and the basal ganglia is prominent at rest but disappears before and during movement execution (Brown, 2007; van Wijk et al., 2017; Cao et al., 2024). These deeper brain structures include the basal ganglia (Brown, 2007; Talakoub et al., 2016; van Wijk et al., 2017; Cao et al., 2024) and the thalamus (Paradiso et al., 2004). This subcortical-cortical beta-band synchrony may act as a generalised, network-wide “break” or gate. While beta-band coupling continuously maintains a stable inhibitory state, a functional reduction of the coupling may be a physiological prerequisite across both basal ganglia and cerebellar loops to release downstream pyramidal populations for movement execution.

### Training-related adaptation of cerebellar dynamics

The strength of cerebellar and cortical beta-band ERDs developed differently throughout the training period. This divergence suggests that cerebellar beta-band ERD specifically reflects adaptations underlying the acquisition of context-dependent motor timing. This slow-scaled adaptation likely serves an inductive role in tuning internal timing models across both ocular and manual effectors to shape accurate context-dependent motor timing (Koppen et al., 2026). Across learning, Purkinje cell simple-spike activity undergoes experience-dependent modification, typically decreasing immediately prior to an expected event (Johansson et al., 2016). This progressive suppression of Purkinje cell discharge is driven by long-term depression at parallel fibre to Purkinje cell synapses, induced by error signals conveyed via climbing fibres that emerge across training (Medina and Lisberger, 2008; Rasmussen et al., 2008).

Here, the cerebellar beta-band ERD observed with OPM-MEG may serve as a non-invasive, macroscopic index of these cumulative synaptic adjustments, capturing the progressive emergence of an accurately timed, feedforward disinhibition of the DCN necessary to release the manual motor command. Interestingly, conventional MEG recordings of the human cerebellum found support that cerebellar beta-band oscillations play a role in predicting the sensory timing of stimuli (Andersen and Dalal, 2021, 2024) with its strength related to the temporal regularity of the stimulations prior to the omission (Andersen and Dalal, 2021). Collectively, this reinforces the role of cerebellar beta-band oscillations as a driver of the experience-dependent plastic changes required to orchestrate predictive, context-dependent timing across motor and sensory domains.

### Conclusion

In sum, human cortico-cerebellar beta-band oscillations serve as a dynamic gating mechanism for flexible temporal control. To allow for well-timed movements, the cerebellum supports the motor cortex in planning and releasing temporally accurate motor commands.

## Author Contributions

D.N., and K.K. designed research; C.C.W., J.W., A.K., and K.K. performed research; C.S.Z. analysed data; C.S.Z., and K.K. wrote the first draft of the paper; C.S.Z., O.J., R.C.M., D.N., and K.K. edited the paper; D.N., and K.K. provided the funding.

## Acknowledgements

C.S.Z. and K.K. were supported by the UKRI Future Leaders Fellowship (awarded to K.K.: MR/Y016467/1) and the Centre for Human Brain Health for OPM-MEG access. D.N. was supported by the Netherlands Organization for Scientific Research (Vidi-VI.193.076, Aspasia-015.016.012 and Gravitation DBI2-024.005.022), the AINED organization and the Nationaal Groeifonds (AINED/NGF NGF.1609.241.021). We are grateful to Dr Massimiliano di Luca for providing the air pressuriser for the eyeblink task.

## References

Ahlfors SP, Han J, Belliveau JW, Hämäläinen MS (2010) Sensitivity of MEG and EEG to source orientation. Brain Topography 23:227–232.

Andersen LM, Dalal SS (2021) The cerebellar clock: Predicting and timing somatosensory touch. NeuroImage 238:118202.

Andersen LM, Dalal SS (2024) Detection of threshold-level stimuli modulated by temporal predictions of the cerebellum. eNeuro 11.

Andersen LM, Jerbi K, Dalal SS (2020) Can EEG and MEG detect signals from the human cerebellum? NeuroImage 215:116817.

Baccalá LA, Sameshima K (2001) Partial directed coherence: a new concept in neural structure determination. Biological Cybernetics 84:463–474.

Barone J, Rossiter HE (2021) Understanding the role of sensorimotor beta oscillations. Frontiers in Systems Neuroscience 15:655886.

Bauer M, Stenner M-P, Friston KJ, Dolan RJ (2014) Attentional modulation of alpha/beta and gamma oscillations reflect functionally distinct processes. The Journal of Neuroscience 34:16117–16125.

Bloedel JR, Courville J (1981) Cerebellar afferent systems. Handbook of Physiology 2:735–829.

Boto E, Holmes N, Leggett J, Roberts G, Shah V, Meyer SS, Muñoz LD, Mullinger KJ, Tierney TM, Bestmann S (2018) Moving magnetoencephalography towards real-world applications with a wearable system. Nature 555:657–661.

Brown P (2007) Abnormal oscillatory synchronisation in the motor system leads to impaired movement. Current Opinion in Neurobiology 17:656–664.

Cao C, Litvak V, Zhan S, Liu W, Zhang C, Sun B, Li D, van Wijk BC (2024) Low-beta versus high-beta band cortico-subcortical coherence in movement inhibition and expectation. Neurobiology of Disease 201:106689.

Chabrol FP, Blot A, Mrsic-Flogel TD (2019) Cerebellar contribution to preparatory activity in motor neocortex. Neuron 103:506–519. e504.

Deverett B, Koay SA, Oostland M, Wang SS (2018) Cerebellar involvement in an evidence-accumulation decision-making task. eLife 7:e36781.

Diedrichsen J, Zotow E (2015) Surface-based display of volume-averaged cerebellar imaging data. PLoS One 10:e0133402.

Engel AK, Fries P (2010) Beta-band oscillations—signalling the status quo? Current Opinion in Neurobiology 20:156–165.

Faes L, Porta A, Nollo G (2009) Surrogate data approaches to assess the significance of directed coherence: application to EEG activity propagation. In: 2009 Annual International Conference of the IEEE Engineering in Medicine and Biology Society, pp 6280–6283: IEEE.

Gao Z, Davis C, Thomas AM, Economo MN, Abrego AM, Svoboda K, De Zeeuw CI, Li N (2018) A cortico-cerebellar loop for motor planning. Nature 563:113–116.

Groß J, Kujala J, Hämäläinen M, Timmermann L, Schnitzler A, Salmelin R (2001) Dynamic imaging of coherent sources: studying neural interactions in the human brain. Proceedings of the National Academy of Sciences 98:694–699.

Groth CL, Singh A, Zhang Q, Berman BD, Narayanan NS (2021) GABAergic modulation in movement related oscillatory activity: A review of the effect pharmacologically and with aging. Tremor and Other Hyperkinetic Movements 11:48.

Hazeltine E, Helmuth LL, Ivry RB (1997) Neural mechanisms of timing. Trends in Cognitive Sciences 1:163–169.

Heiney SA, Wohl MP, Chettih SN, Ruffolo LI, Medina JF (2014) Cerebellar-dependent expression of motor learning during eyeblink conditioning in head-fixed mice. Journal of Neuroscience 34:14845–14853.

Hillebrand A, Barnes GR (2002) A quantitative assessment of the sensitivity of whole-head MEG to activity in the adult human cortex. NeuroImage 16:638–650.

Holmes CJ, Hoge R, Collins L, Woods R, Toga AW, Evans AC (1998) Enhancement of MR images using registration for signal averaging. Journal of Computer Assisted Tomography 22:324–333.

Israely S, Ninou H, Rajchert O, Elmaleh L, Harel R, Mawase F, Kadmon J, Prut Y (2025) Cerebellar output shapes cortical preparatory activity during motor adaptation. Nature Communications 16:2574.

Johansson F, Hesslow G, Medina JF (2016) Mechanisms for motor timing in the cerebellar cortex. Current Opinion in Behavioral Sciences 8:53–59.

King M, Hernandez-Castillo CR, Poldrack RA, Ivry RB, Diedrichsen J (2019) Functional boundaries in the human cerebellum revealed by a multi-domain task battery. Nature Neuroscience 22:1371–1378.

Koppen J, Klinkhamer I, Runge M, Bayones L, Narain D (2026) Neural circuits encode prior knowledge of temporal statistics. Nature Neuroscience:1–10.

Kornysheva K (2016) Encoding temporal features of skilled movements—what, whether and how? Progress in Motor Control: Theories and Translations:35–54.

Kujala J, Pammer K, Cornelissen P, Roebroeck A, Formisano E, Salmelin R (2007) Phase coupling in a cerebro-cerebellar network at 8–13 Hz during reading. Cerebral Cortex 17:1476–1485.

Lin C-HS, Tierney TM, Mellor S, O’Neill GC, Bestmann S, Barnes GR, Miall RC (2025) Early insights into eyeblink conditioning using optically pumped magnetometer-based MEG. Frontiers in Human Neuroscience 19:1638751.

Lin CH, Tierney TM, Holmes N, Boto E, Leggett J, Bestmann S, Bowtell R, Brookes MJ, Barnes GR, Miall RC (2019) Using optically pumped magnetometers to measure magnetoencephalographic signals in the human cerebellum. The Journal of Physiology 597:4309–4324.

Mangili L, Wissing C, Narain D (2025) Fast implicit and slow explicit learning of temporal context. Scientific Reports 15:16343.

Maris E, Oostenveld R (2007) Nonparametric statistical testing of EEG-and MEG-data. Journal of Neuroscience Methods 164:177–190.

Medina JF, Lisberger SG (2008) Links from complex spikes to local plasticity and motor learning in the cerebellum of awake-behaving monkeys. Nature Neuroscience 11:1185–1192.

Medina JF, Nores WL, Ohyama T, Mauk MD (2000) Mechanisms of cerebellar learning suggested by eyelid conditioning. Current Opinion in Neurobiology 10:717–724.

Merchant H, Pérez O, Zarco W, Gámez J (2013) Interval tuning in the primate medial premotor cortex as a general timing mechanism. Journal of Neuroscience 33:9082–9096.

Nobre AC, Correa A, Coull JT (2007) The hazards of time. Current Opinion in Neurobiology 17:465–470.

Nolte G (2003) The magnetic lead field theorem in the quasi-static approximation and its use for magnetoencephalography forward calculation in realistic volume conductors. Physics in Medicine & Biology 48:3637.

Nunez PL, Srinivasan R (2006) A theoretical basis for standing and traveling brain waves measured with human EEG with implications for an integrated consciousness. Clinical Neurophysiology 117:2424–2435.

Oldfield RC (1971) The assessment and analysis of handedness: the Edinburgh inventory. Neuropsychologia 9:97–113.

Oostenveld R, Fries P, Maris E, Schoffelen J-M (2011) FieldTrip: open source software for advanced analysis of MEG, EEG, and invasive electrophysiological data. Computational Intelligence and Neuroscience 2011:156869.

Paradiso G, Cunic D, Saint-Cyr JA, Hoque T, Lozano AM, Lang AE, Chen R (2004) Involvement of human thalamus in the preparation of self-paced movement. Brain 127:2717–2731.

Pfurtscheller G, Da Silva FL (1999) Event-related EEG/MEG synchronization and desynchronization: basic principles. Clinical Neurophysiology 110:1842–1857.

Rasmussen A, Jirenhed D-A, Hesslow G (2008) Simple and complex spike firing patterns in Purkinje cells during classical conditioning. The Cerebellum 7:563–566.

Remington ED, Egger SW, Narain D, Wang J, Jazayeri M (2018) A dynamical systems perspective on flexible motor timing. Trends in Cognitive Sciences 22:938–952.

Roos S, Hämäläinen M, Iivanainen J (2026) Efficacy of Optically Pumped Magnetometers in Detecting Activity From the Cerebellar Cortex. Human Brain Mapping 47:e70514.

Rorden C (2025) MRIcroGL: voxel-based visualization for neuroimaging. Nature Methods 22:1613–1614.

Samuelsson JG, Sundaram P, Khan S, Sereno MI, Hämäläinen MS (2020) Detectability of cerebellar activity with magnetoencephalography and electroencephalography. Human Brain Mapping 41:2357–2372.

Sinha N, Israely S, Ben Harosh O, Harel R, Dewald JP, Prut Y (2025a) Disentangling acute motor deficits and adaptive responses evoked by the loss of cerebellar output. eLife 14:RP105152.

Sinha N, Yao J, Harosh OB, Harel R, Dewald JP, Prut Y (2025b) Cerebellar Output to the Motor Cortex Facilitates Beta Band Suppression During Movement Preparation. In: Summer School on Neurorehabilitation, pp 160–164: Springer.

Stine GM, Jazayeri M (2025) Control principles of neural dynamics revealed by the neurobiology of timing. Annual Review of Neuroscience 48:43–63.

Talakoub O, Neagu B, Udupa K, Tsang E, Chen R, Popovic MR, Wong W (2016) Time-course of coherence in the human basal ganglia during voluntary movements. Scientific Reports 6:34930.

Ten Brinke MM, Boele H-J, Spanke JK, Potters J-W, Kornysheva K, Wulff P, IJpelaar AC, Koekkoek SK, De Zeeuw CI (2015) Evolving models of pavlovian conditioning: cerebellar cortical dynamics in awake behaving mice. Cell Reports 13:1977–1988.

Ten Brinke MM, Heiney SA, Wang X, Proietti-Onori M, Boele H-J, Bakermans J, Medina JF, Gao Z, De Zeeuw CI (2017) Dynamic modulation of activity in cerebellar nuclei neurons during pavlovian eyeblink conditioning in mice. eLife 6:e28132.

Tesche CD, Karhu JJ (2000) Anticipatory cerebellar responses during somatosensory omission in man. Human Brain Mapping 9:119–142.

Tzourio-Mazoyer N, Landeau B, Papathanassiou D, Crivello F, Etard O, Delcroix N, Mazoyer B, Joliot M (2002) Automated anatomical labeling of activations in SPM using a macroscopic anatomical parcellation of the MNI MRI single-subject brain. NeuroImage 15:273–289.

van Wijk BC, Neumann W-J, Schneider G-H, Sander TH, Litvak V, Kühn AA (2017) Low-beta cortico-pallidal coherence decreases during movement and correlates with overall reaction time. NeuroImage 159:1–8.

Wagner MJ, Kim TH, Kadmon J, Nguyen ND, Ganguli S, Schnitzer MJ, Luo L (2019) Shared cortex-cerebellum dynamics in the execution and learning of a motor task. Cell 177:669–682. e624.

Wang J, Narain D, Hosseini EA, Jazayeri M (2018) Flexible timing by temporal scaling of cortical responses. Nature Neuroscience 21:102–110.

West TO, Spedden ME, O’Neill GC, Bestmann S, Barnes G, Farmer SF, Cagnan H (2025) The Cerebellum Implements an Oscillatory Forward Model for Accurate Motor Timing. bioRxiv:2025.2011. 2011.687835.

Westner BU, Dalal SS, Gramfort A, Litvak V, Mosher JC, Oostenveld R, Schoffelen J-M (2022) A unified view on beamformers for M/EEG source reconstruction. NeuroImage 246:118789.

