## Supplementary_material for "Cortico-cerebellar beta-band dynamics predict flexible motor timing"

### *Supplementary figures*

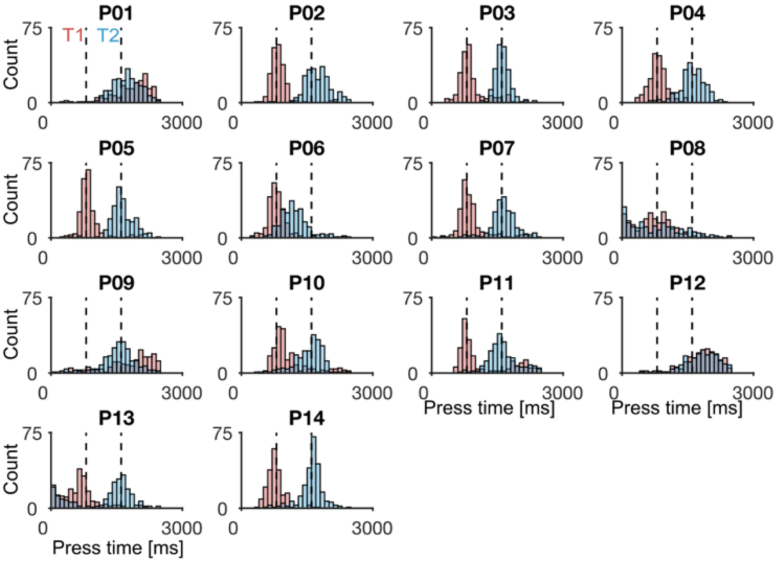


***Figure S1. Participant- and context-dependent motor timings.*** *Histograms are depicted with a bin width of 100 ms. Context-dependent distributions are shown in red (T1) and blue (T2). Graph titles indicate participant coding (e.g., P01: participant 1). Black vertical graphs indicate the target timings at T1 and T2.*

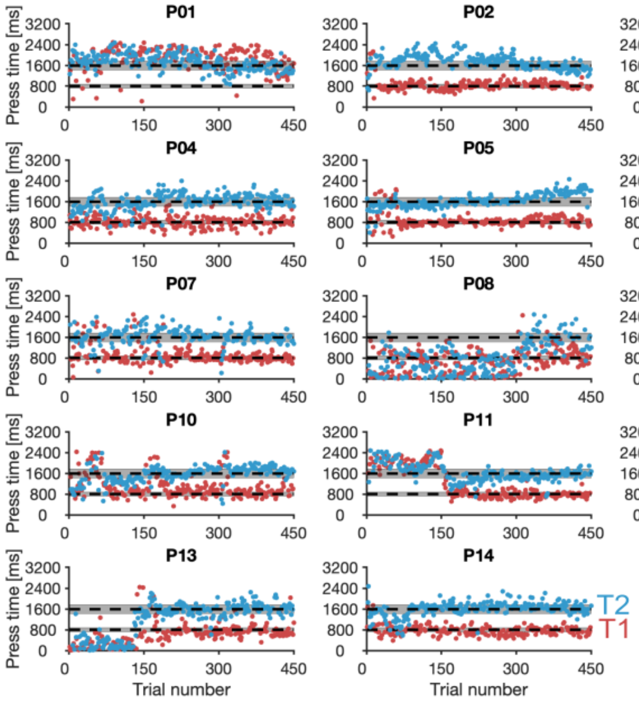


***Figure S2. Participant- and context-dependent manual timings over the course of the experiment.*** *Vertical grey patches represent the respective omission windows and black horizontal graphs indicate the target timings at T1 and T2. Graph titles indicate participant coding (e.g., P01: participant 1).*

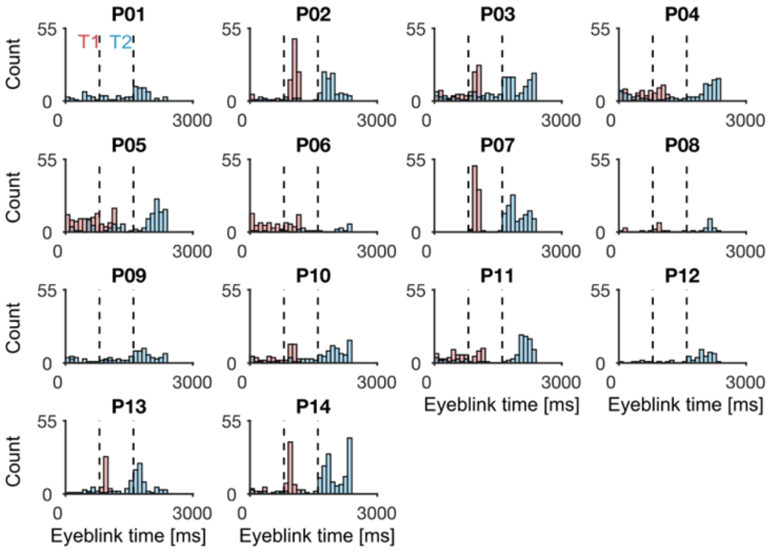


***Figure S3. Participant- and context-dependent conditioned eyeblink timings.*** *Histograms are depicted with a bin width of 100 ms. Context-dependent distributions are shown in red (T1) and blue (T2). Graph titles indicate participant coding (e.g., P01: participant 1). Black vertical graphs indicate the target timings at T1 and T2.*

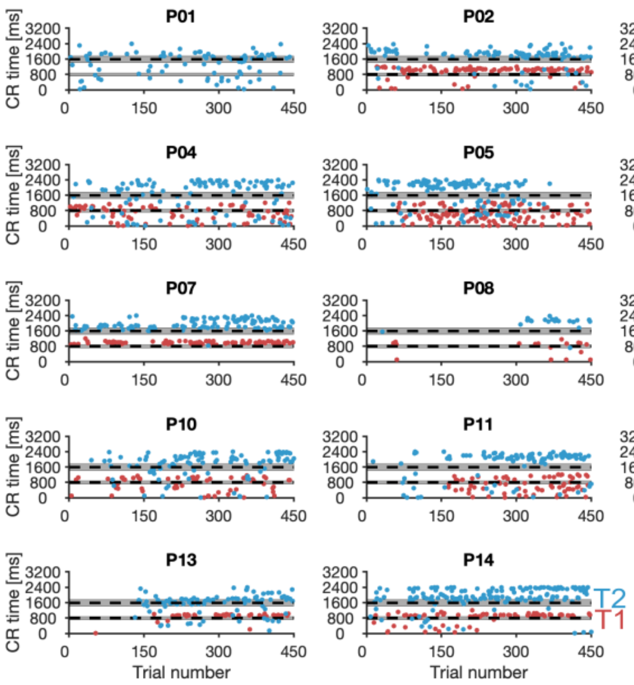


***Figure S4. Participant- and context-dependent conditioned eyeblink timings over the course of the experiment.*** *Vertical grey patches represent the respective omission windows and black horizontal graphs indicate the target timings at T1 and T2. Graph titles indicate participant coding (e.g., P01: participant 1).*

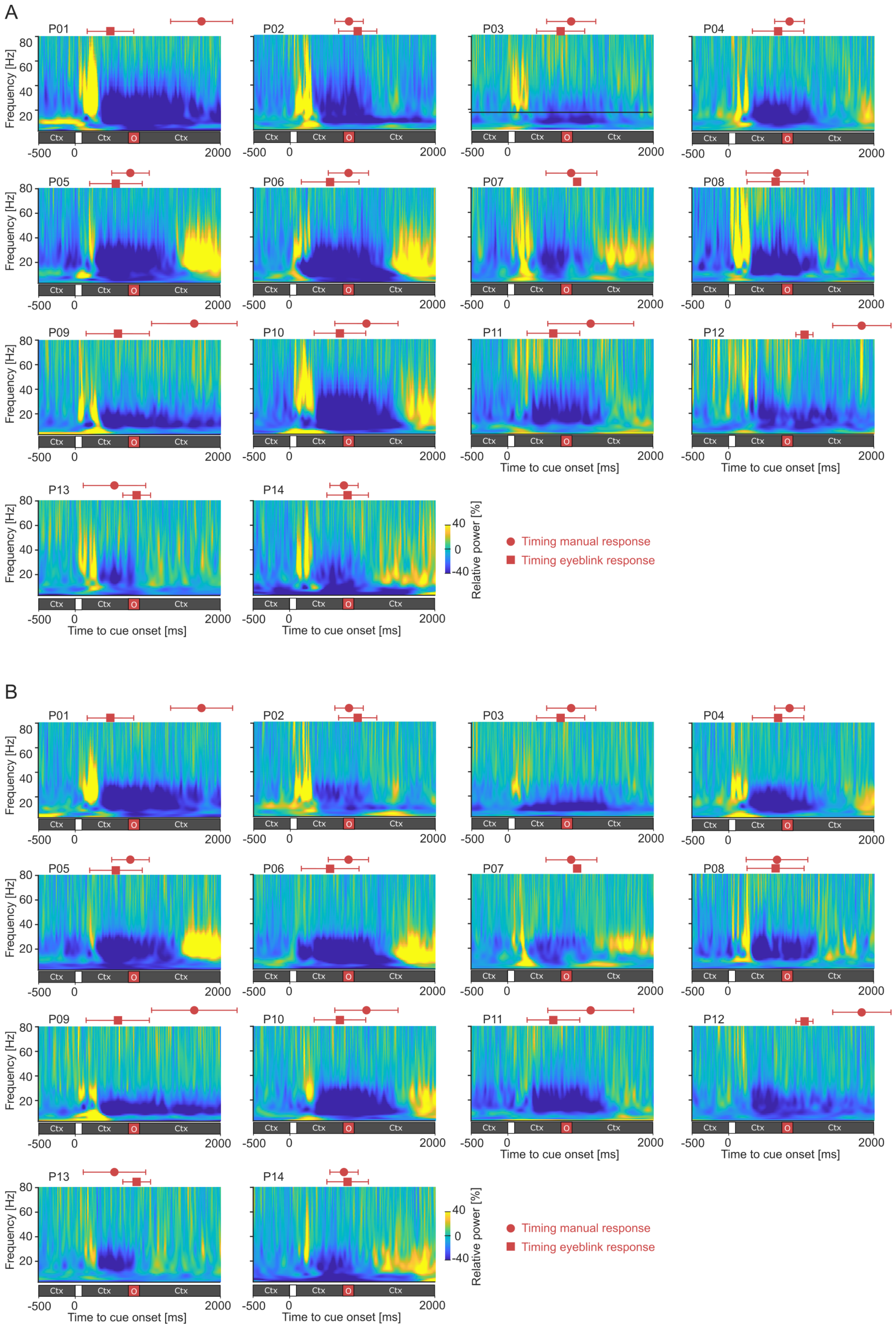
A
B
P01
P02
P03
P04
P05
P06
P07
P08
P09
P10
P11
P12
P13
P14
Frequency [Hz]
80
60
40
20
Ctx
O
-500
0
2000
Time to cue onset [ms]
Relative power [%]
40
0
-40
Timing manual response
Timing eyeblink response

***Figure S5.*** *Participant-specific time-frequency-power representations of left cerebellar sensor* ***A)*** *L115 and* ***B)*** *R115 for T1. Red circles and squares are the mean times of the manual and conditioned eyeblink response events (error bars are standard deviations over trials). Bar below the time-frequency representation indicate the timings within the trial. The end of the brief flash at 0 sec marked the start of the trial and of the context starting to move. Graph titles indicate participant coding (e.g., P01: participant 1). Ctx: context; O: omission window.*

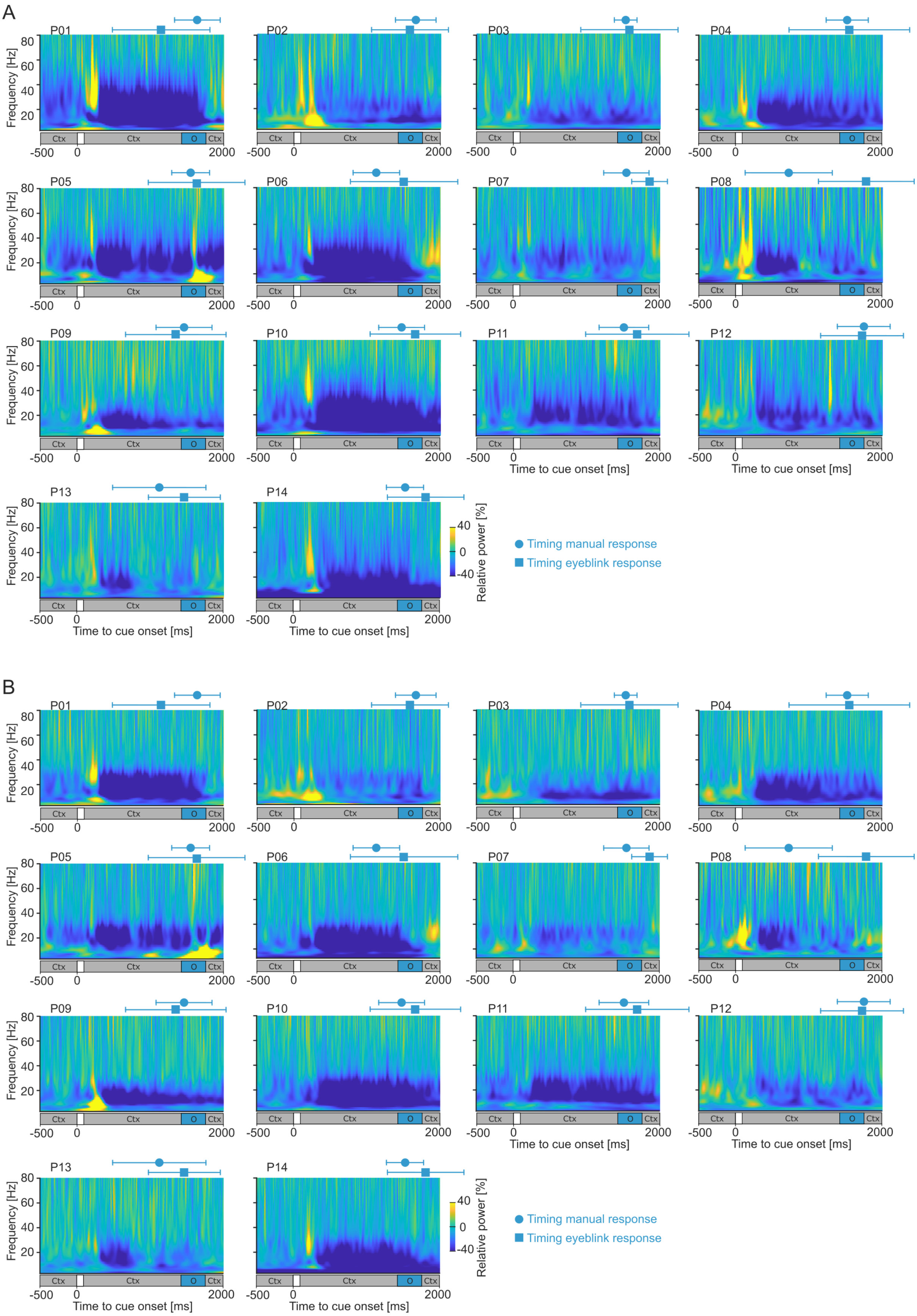


***Figure S6.*** *Participant-specific time-frequency-power representations of left cerebellar sensor* ***A)*** *L115 and* ***B)*** *R115 for T2. Blue circles and squares are the mean times of the manual and conditioned*

*eyeblink response events (error bars are standard deviations over trials). Bar below the time-frequency representation indicate the timings within the trial. The end of the brief flash at 0 sec marked the start of the trial and of the context starting to move. Graph titles indicate participant coding (e.g., P01: participant 1). Ctx: context; O: omission window.*

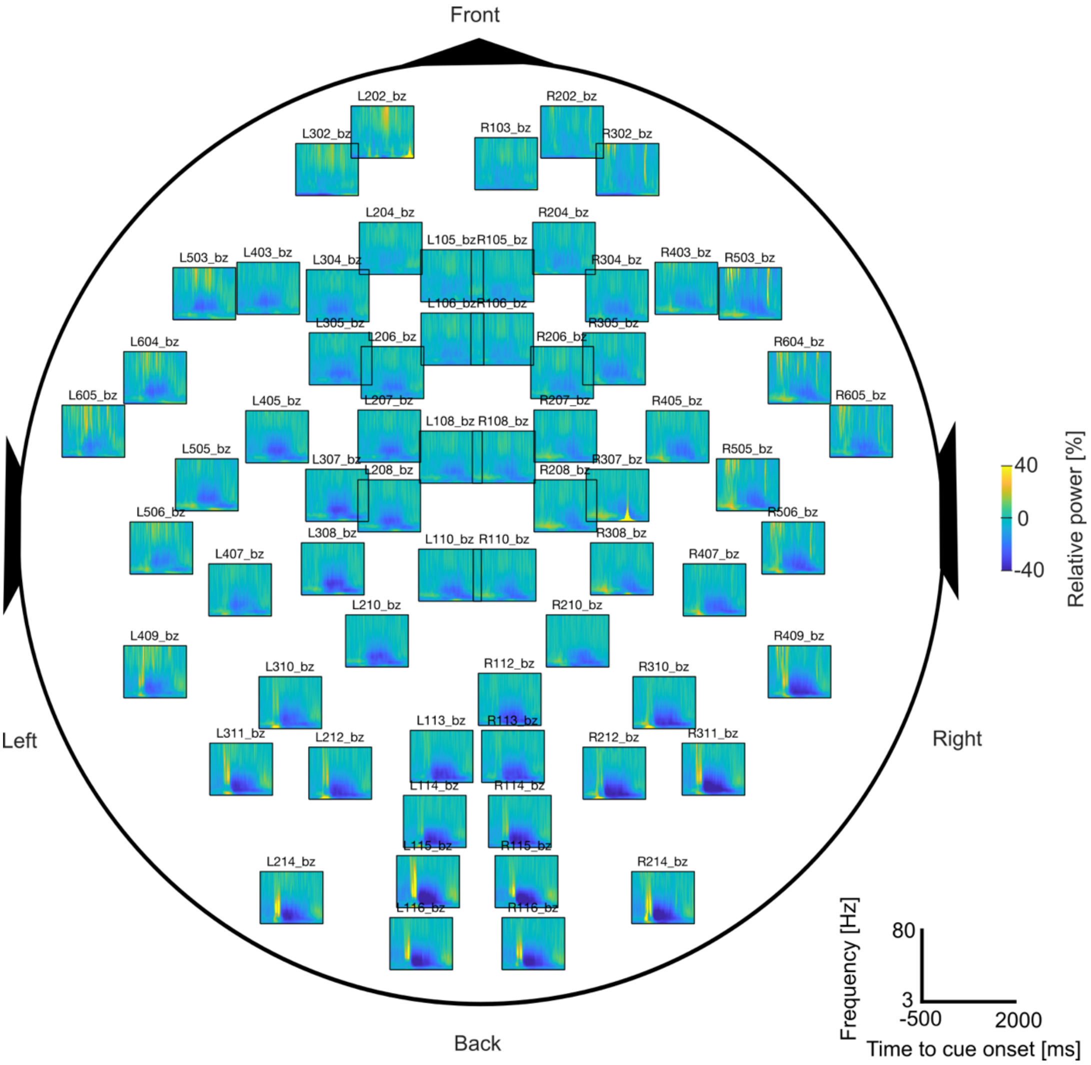


***Figure S7. Time-frequency-power representations for T1 for all sensors.*** *Representations depict averages over all participants.*

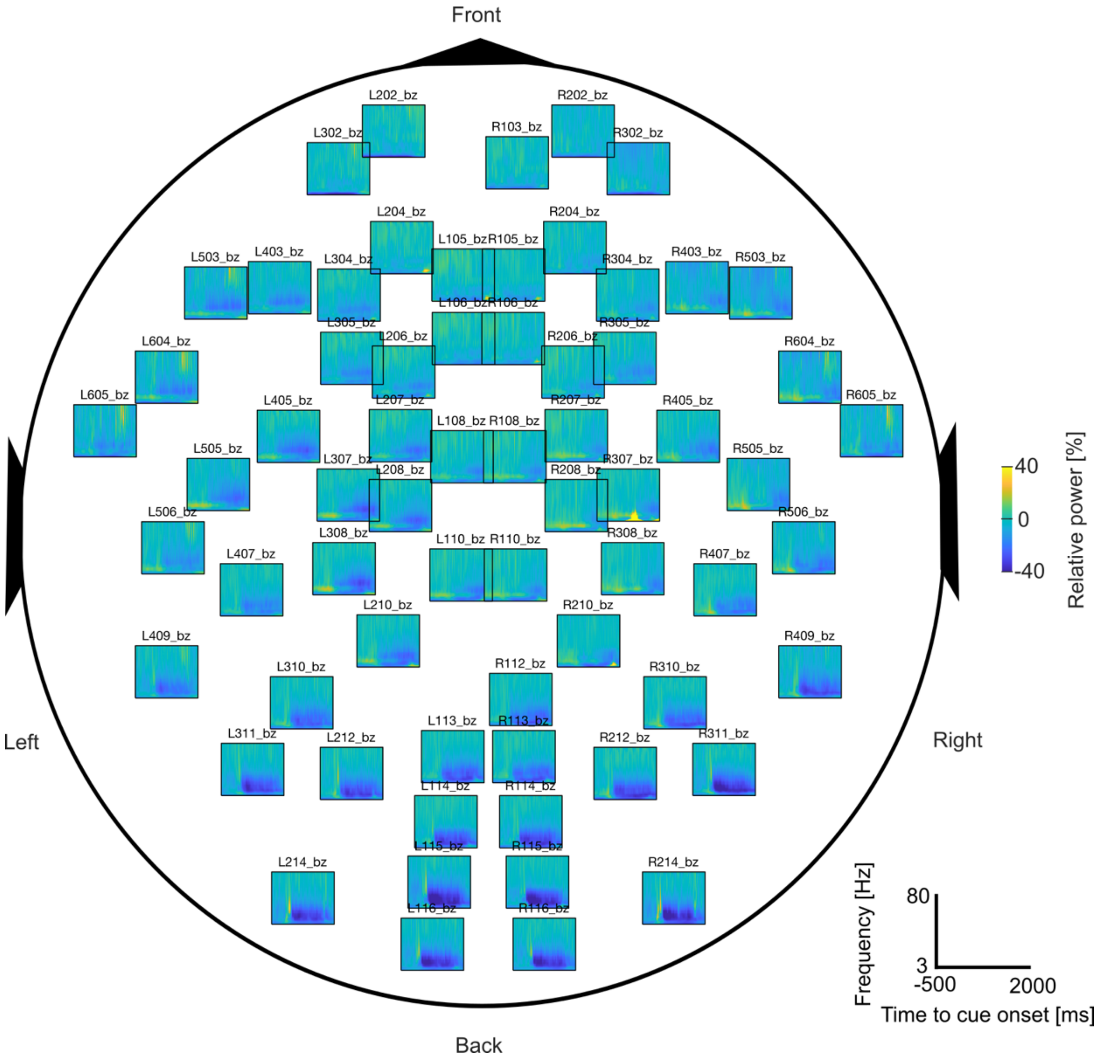


***Figure S8. Time-frequency-power representations for T2 for all sensors.*** *Representations depict averages over all participants.*

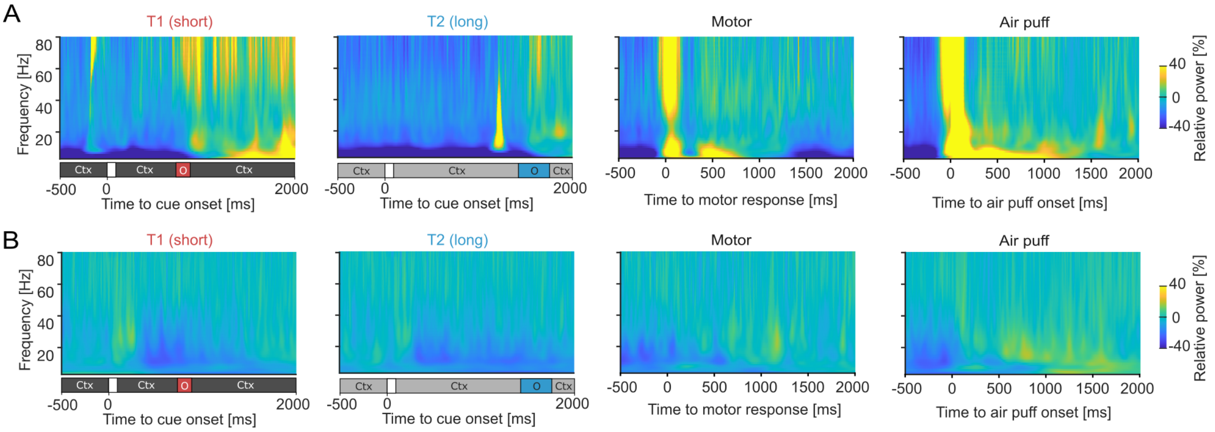


***Figure S9. Time-frequency-power representations of the A) magneto-oculography of left eye and B) magnetomyography over the right trapezius muscle (R117).*** *Bar below the time-frequency representation indicate the timings within the trial. The end of the brief flash at 0 sec marked the start of the trial and of the context starting to move. Ctx: context; O: omission window.*

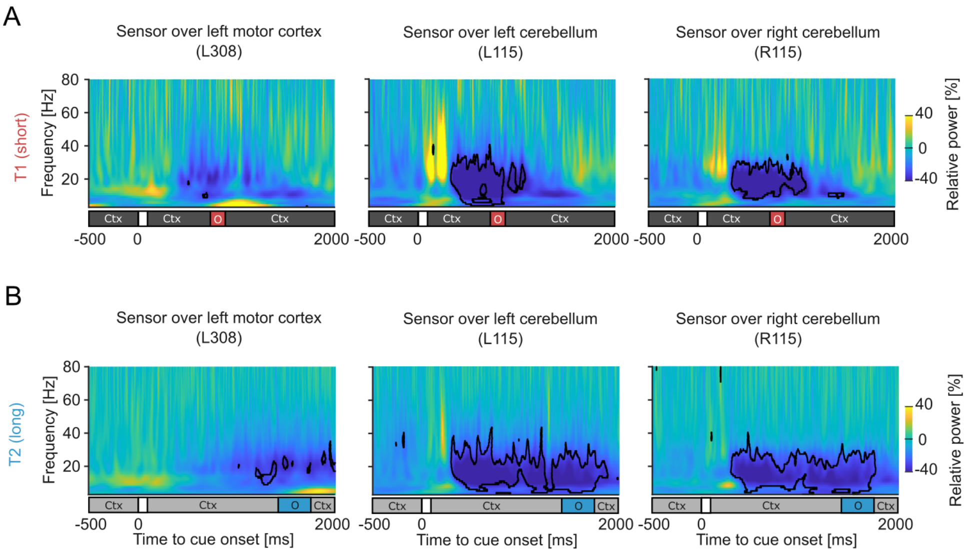


***Figure S10. Time-frequency-power representations of sensors over the left motor cortex and bilateral cerebellum for correct manual trials.*** *Time-frequency-power representations of the T1 for sensors L308 (left motor cortex), L115 (left cerebellum) and R115 (right cerebellum). Bar below the time-frequency representation indicate the timings within the trial. The end of the brief flash at 0 sec marked the start of the trial and of the context starting to move. Black outlines indicate statistical significance (p: <0.05; false discovery rate). Ctx: context; O: omission window.*

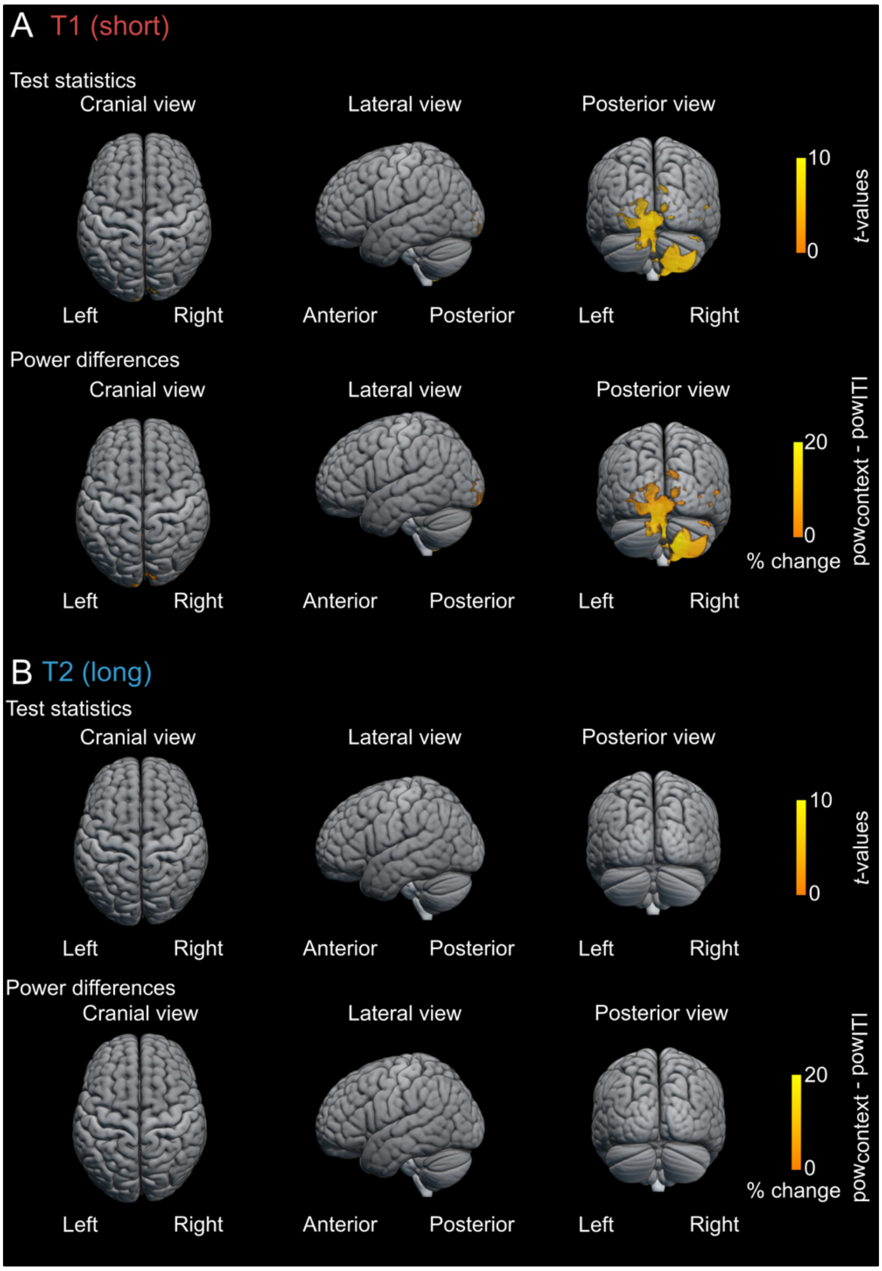


***Figure S11. Source localisations of test statistics, contrasting context- and inter-trial-interval periods.*** *Source localisations aimed to localise the gamma-band synchronisation (frequency range: 30-80 Hz; time window: 0-300 ms; p: <0.05 false discovery rate) for* ***A)*** *T1 and* ***B)*** *T2. Statistical masking is done using the alpha-level thresholds.*

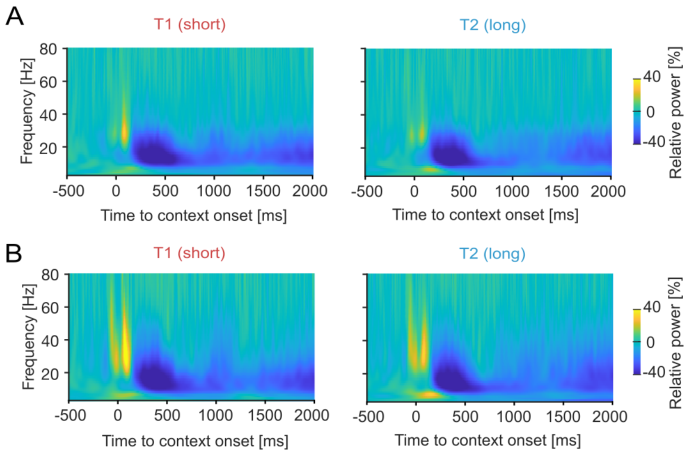


***Figure S12. Sensor-level cerebellar responses time-locked to the (static) contexts.*** *Cerebellar responses are provided for sensors over the **A)** left (L115) and **B)** right (R115) cerebellum.*

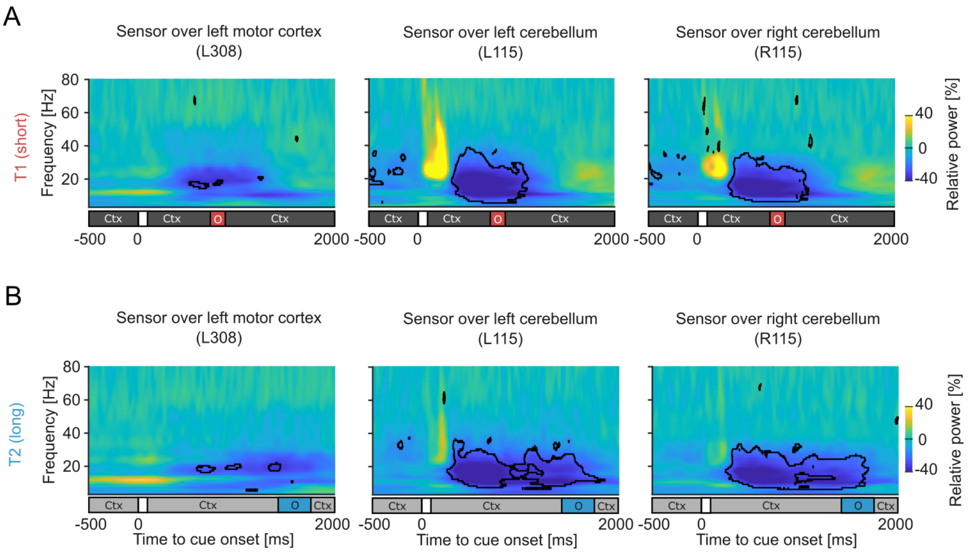


***Figure S13. Time-frequency-power representations of sensors over the left motor cortex and bilateral cerebellum with Hanning windows with a length of 7 cycles of the frequency of interest.*** *Time-frequency-power representations of the T1 for sensors L308 (left motor cortex), L115 (left cerebellum) and R115 (right cerebellum). Bar below the time-frequency representation indicate the timings within the trial. The end of the brief flash at 0 sec marked the start of the trial and of the context starting to move. Black outlines indicate statistical significance (p: <0.05; false discovery rate). Ctx: context; O: omission window.*

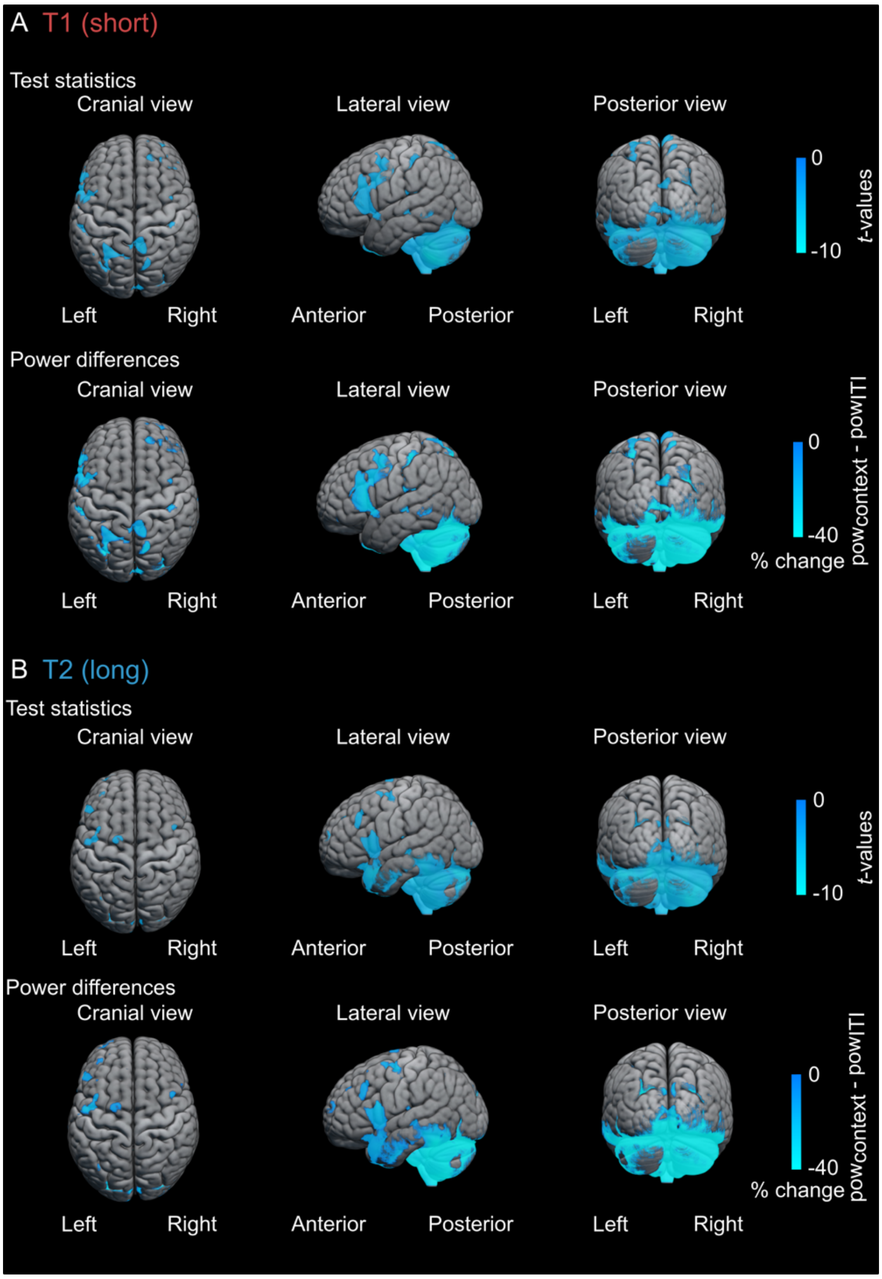


***Figure S14. Spatial localisations for A) T1 and B) T2.*** *Source localisations are constrained to alpha-band frequencies (frequency range: 8-12 Hz) and context (time window T1: 400 to 900 ms; time window T2: 400 to 1,700 ms). Test statistics and power differences are masked by the statistical contrasts between context and inter-trial interval (p: <0.001 false discovery rate).*

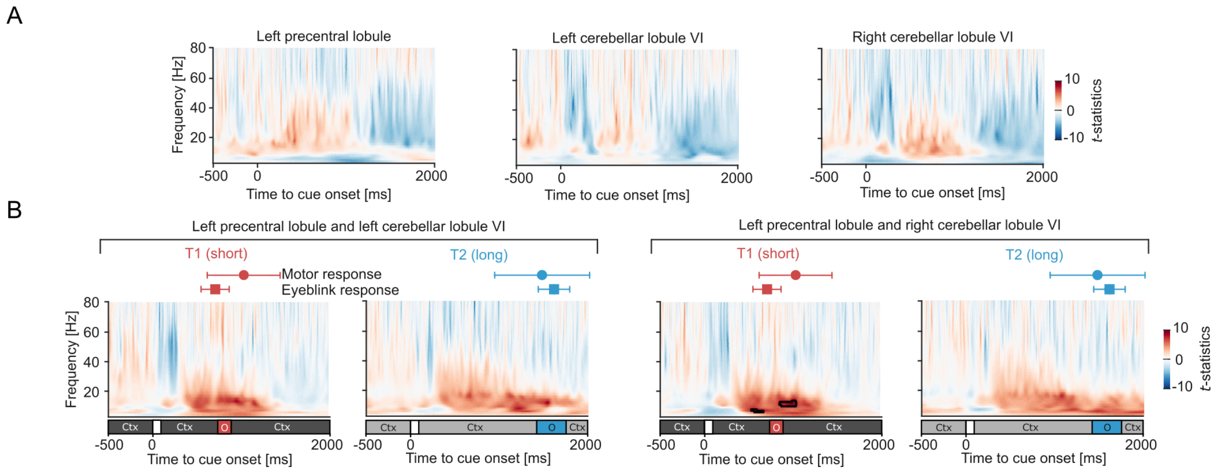


***Figure S15. A)*** *Cross-context statistical comparisons (p: $<0.05$; FDR corrected) between T1 and T2 for the left precentral lobule and bilateral cerebellar lobules VI. Positive and negative t-statistics indicate mean differences for $T1 < T2$ and $T1 > T2$, respectively.* ***B)*** *Cross-region statistical comparisons (p: $<0.05$; FDR corrected) between left precentral lobule and bilateral cerebellar lobules VI. Positive and negative t-statistics indicate cerebellum > precentral lobule and cerebellum < precentral lobule, respectively. Ctx: context; O: omission window.*

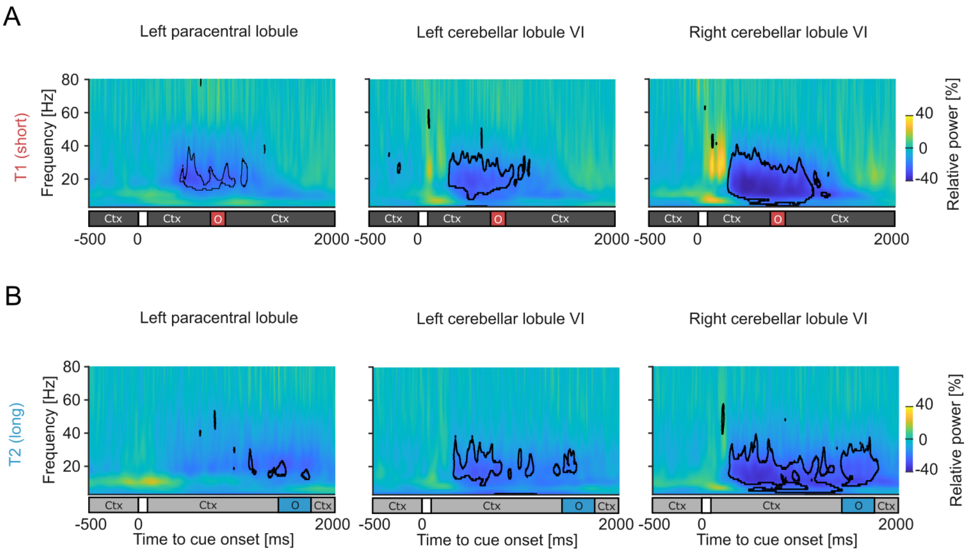


***Figure S16. Source-reconstructed time-frequency-power representations using mean spatial filters.*** *Time-frequency-power representations of T1 and T2 for the left paracentral lobule and bilateral cerebellar lobules VI. Black outlines indicate statistical significance (p: <0.05; false discovery rate). Bar below the time-frequency representation indicates the timings within the trial. The static context was followed by a brief flash of 100 ms after which the context started moving. Ctx: context; O: omission window.*

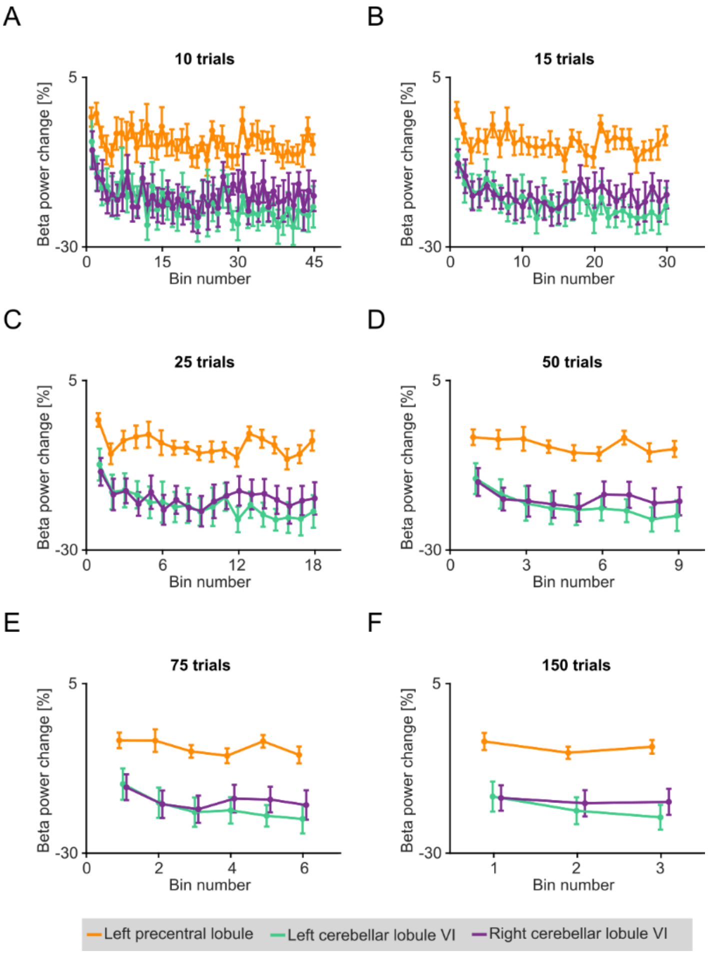


***Figure S17. Training-related beta-band power changes for different bin sizes.*** *Changes are expressed relative to the inter-trial interval and pooled into* ***A)*** *10-trial,* ***B)*** *15-trial,* ***C)*** *25-trial,* ***D)*** *50-trial,* ***E)*** *75-trial, and* ***F)*** *150-trial bins. Bars and error bars represent group means and standard errors of the mean over participants.*

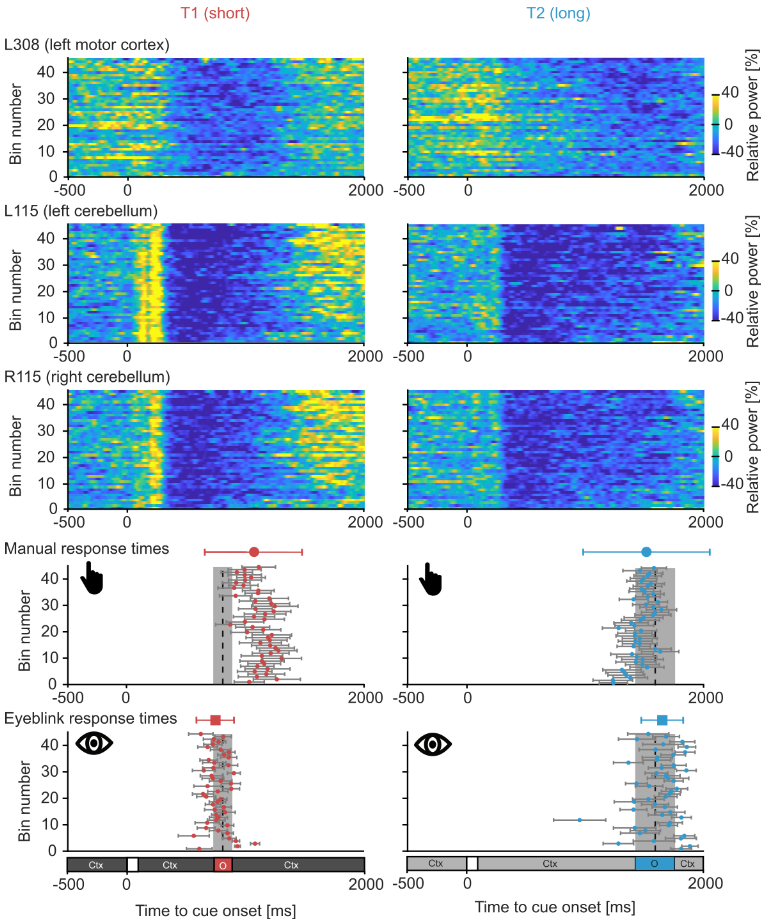


***Figure S18. Training-related beta-band power changes at sensor level and its relation to manual and eyeblink responses. A)*** *Time-resolved power changes for sensors over the left motor cortex (L308) and left and right cerebellar (L115 and R115, respectively) areas for both contexts aligned to the timings of the manual and conditioned eyeblink responses. Power is averaged over the frequencies of 13 and 30 Hz (comprising beta-band frequencies) and binned over every 5 trials. Manual response and conditioned eyeblink times are the mean and standard error of the mean. Bar below the conditioned eyeblink response times indicates the timings within the trial. The static context was followed by a brief flash of 100 ms after which the context started moving. Ctx: context; O: omission window.*

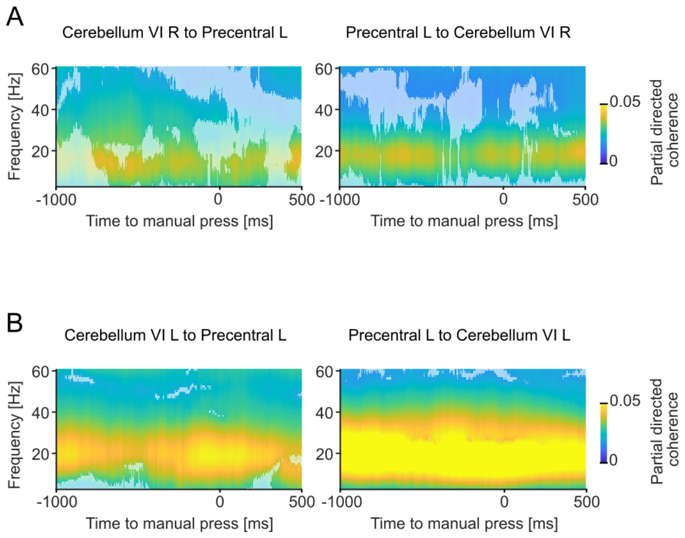


***Figure S19. Cortico-cerebellar coupling time-locked to the manual response timing.*** *Partial directed coherence between the left paracentral lobule and* ***A)*** *right and* ***B)*** *left cerebellar lobule VI. Opaque samples indicate statistical significance relative to surrogate coupling (p: <0.05; false discovery rate). Note that the derived partial directed coherence only comprises correct trials (i.e., trials in which participants correctly pressed a button within the omission window).*

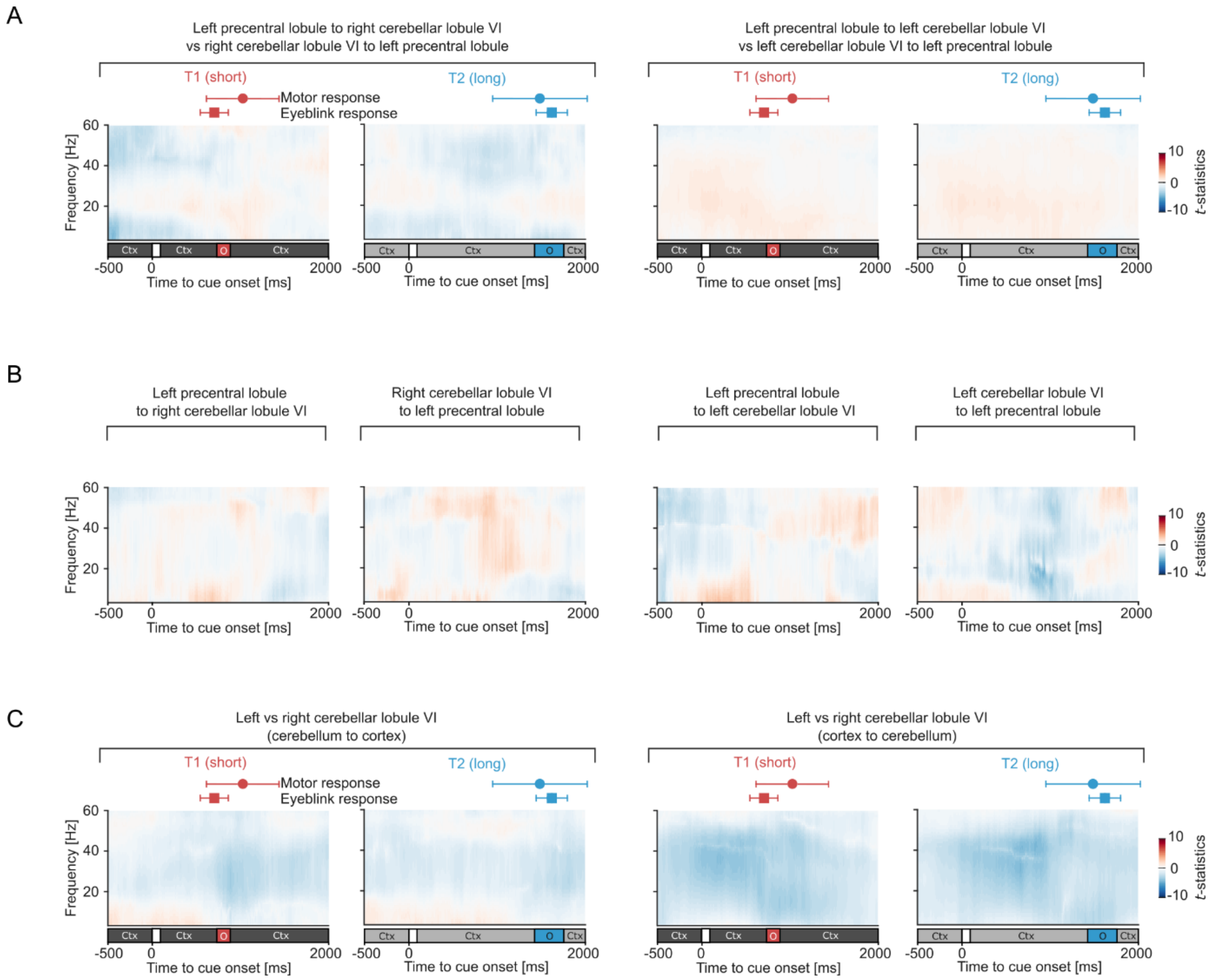


***Figure S20. Statistical evaluations of cortico-cerebellar coupling across directionality, contexts, and cerebellar sides. A)*** *Cross-directionality statistical comparisons (p: >0.05; FDR corrected) for both contexts and cerebellar hemispheres. Positive and negative t-statistics indicate mean differences for cortex to cerebellum < cerebellum to cortex, and cortex to cerebellum > cerebellum to cortex, respectively. Ctx: context; O: omission window.* ***B)*** *Cross-context statistical comparisons (p: >0.05; FDR corrected) for both directionalities and cerebellar hemispheres. Positive and negative t-statistics indicate mean differences for T2 < T1 and T2 > T1, respectively.* ***C)*** *Cross-hemisphere statistical comparisons (p: >0.05; FDR corrected) for both directionalities and contexts. Positive and negative t-statistics indicate mean differences for left cerebellum < right cerebellum, and left cerebellum > right cerebellum, respectively.*

***Supplementary tables***

| **Region** | **Maximal change [%]** | **Mean change [%]** | **Location maximum [mm]** | ***P*-value** | **Number of voxels** |
|---|---|---|---|---|---|
| **Cerebellar lobule VI right** | -37.36 | -32.12 | [30, -75, -20] | 7e-6 | 117 |
| **Cerebellar crus I right** | -36.92 | -31.91 | [20, -80, -25] | 5e-6 | 155 |
| **Cerebellar crus II right** | -36.46 | -31.47 | [20, -80, -35] | 4e-6 | 141 |
| **Cerebellar lobule VI left** | -36.53 | -31.13 | [-20, -70, -25] | 8e-7 | 106 |
| **Cerebellar crus II left** | -35.43 | -31.09 | [-20, -80, -35] | 3e-7 | 112 |
| **Cerebellar lobule VIII right** | -35.46 | -31.01 | [20, -70, -40] | 4e-6 | 149 |
| **Cerebellar lobule VIIb right** | -35.17 | -30.90 | [20, -75, -45] | 4e-6 | 33 |
| **Cerebellar crus I left** | -36.92 | -29.76 | [-20, -75, -30] | 8e-7 | 172 |
| **Cerebellar lobule VIII left** | -34.63 | -29.53 | [-15, -70, -45] | 9e-6 | 122 |
| **Cerebellar lobule IX left** | -31.68 | -28.89 | [-10, -60, -50] | 2e-5 | 50 |
| **Cerebellar lobule IX right** | -30.92 | -28.58 | [10, -60, -45] | 4e-6 | 49 |
| **Cerebellar lobule VIIb left** | -34.41 | -27.43 | [-15, -75, -50] | 5e-6 | 42 |
| **Cerebellar lobule IV/V right** | -32.17 | -26.69 | [10, -60, -10] | 8e-6 | 53 |
| **Cerebellar lobule IV/V left** | -31.58 | -25.87 | [-10, -60, -10] | 5e-6 | 77 |
| **Cerebellar lobule X right** | -26.16 | -24.58 | [20, -40, -45] | 2e-6 | 10 |
| **Cerebellar lobule X left** | -26.20 | -23.41 | [-15, -40, -45] | 2e-5 | 6 |
| **Cerebellar lobule III right** | -26.15 | -23.12 | [10, -45, -25] | 2e-6 | 12 |
| **Cerebellar lobule III left** | -25.38 | -23.07 | [-5, -45, -15] | 6e-6 | 10 |

| | | | | | |
|---|---|---|---|---|---|
| **Precentral lobule left** | -28.93 | -18.92 | [-50, -5, 30] | 3e-5 | 6 |

***Table S1. Characteristics of beta-band desynchronisation in cerebellar and cortical motor regions of interest.*** *Mean and maximal changes are expressed relative to power at inter-trial interval. Changes are provided as a combined estimate for the bilateral cerebellar areas as these values were used for region of interest selection. Changes were taken after masking using the statistical contrasts between context and inter-trial interval (p: <0.001 false discovery rate). Locations and p-values are hemisphere-specific and belong to the voxel displaying the maximal change. Locations are in MNI coordinates.*
